# Bounded rational decision-making models suggest capacity-limited concurrent motor planning in human posterior parietal and frontal cortex

**DOI:** 10.1101/2021.07.13.452019

**Authors:** Sonja Schach, Axel Lindner, Daniel Alexander Braun

## Abstract

While traditional theories of sensorimotor processing have often assumed a serial decision-making pipeline, more recent approaches have suggested that multiple actions may be planned concurrently and vie for execution. Evidence for the latter almost exclusively stems from electrophysiological studies in posterior parietal and premotor cortex of monkeys. Here we study concurrent prospective motor planning in humans by recording functional magnetic resonance imaging (fMRI) during a delayed response task engaging movement sequences towards multiple potential targets. We find that also in human posterior parietal and premotor cortex delay activity modulates both with sequence complexity and the number of potential targets. We tested the hypothesis that this modulation is best explained by concurrent prospective planning as opposed to the mere maintenance of potential targets in memory. We devise a bounded rationality model with information constraints that optimally assigns information resources for planning and memory for this task and determine predicted information profiles according to the two hypotheses. When regressing delay activity on these model predictions, we find that the concurrent prospective planning strategy provides a significantly better explanation of the fMRI-signal modulations. Moreover, we find that concurrent prospective planning is more costly and thus limited for most subjects, as expressed by the best fitting information capacities. We conclude that bounded rational decision-making models allow relating both behavior and neural representations to utilitarian task descriptions based on bounded optimal information-processing assumptions.

**Author summary:** When the future is uncertain, it can be beneficial to concurrently plan several action possibilities in advance. Electrophysiological research found evidence in monkeys that brain regions in posterior parietal and promotor cortex are indeed capable of planning several actions in parallel. We now used fMRI to study brain activity in these brain regions in humans. For our analyses we applied bounded rationality models that optimally assign information resources to fMRI activity in a complex motor planning task. We find that theoretical information costs of concurrent prospective planning explained fMRI activity profiles significantly better than assuming alternative memory-based strategies. Moreover, exploiting the model allowed us to quantify the individual capacity limit for concurrent planning and to relate these individual limits to both subjects’ behavior and to their neural representations of planning.

## 1 Introduction

Traditional theories of sensorimotor processing often consider a sequential pipeline from perception to action, where in between the cognitive system makes a decision that is subsequently implemented by the motor system that plans and executes the corresponding action [1, 2, 3, 4, 5, 6]. In this view perceptual, cognitive, and motor representations do not overlap and are separated in time. Thus, it is often assumed that the value of different alternatives is computed and used for decision-making in prefrontal regions, which is then subsequently translated into action planning in premotor regions, including parietal, precentral, and subcortical regions [1]. In contrast, more recent frameworks, like the affordance competition hypothesis [7, 8, 9], have put forward the notion that actions might be planned simultaneously in several frontoparietal brain areas and vie for execution, whenever opportunities for multiple potential actions arise at any one time.

In line with these more recent frameworks, neurophysiological studies in monkeys have found increasing evidence, that multiple potential actions are planned in parallel and that competitive potential plans are simultaneously represented in posterior parietal cortex (PPC) [10] and dorsal premotor cortex (PMd) [11, 8], although some studies have also credibly argued against such evidence [12, 13]. Moreover, it has been suggested that neurons that are involved in motor planning are often also involved in the decision-making process [14, 8]. In how far particular decisions are accompanied by more integrated or more serial processing may also depend on the task specification, as shown by Cui et al. [2] for spatial and non-spatial decisions. In particular, their results show that different areas in the posterior parietal cortex encode both potential and selected reach plans (parietal reach region) or encode only selected reach plans (dorsal area 5), suggesting a parallel visuomotor cortical circuitry for spatial effector decisions. Some studies have even suggested that parallel planning can take place after movement onset in a way that is effecting movement kinematics and variability during reaching [15, 16, 17].

While most evidence for *parallel* prospective planning towards alternative targets has been acquired in electrophysiological studies of monkey posterior parietal and premotor cortex, fMRI-studies in humans so far have mostly revealed a contribution of these areas to the prospective planning of movements towards single targets (e.g. [18]). A particular challenge when studying parallel processing in fMRI is its limited temporal resolution, so what might seem like parallel multi-target planning on the coarse timescale of fMRI might, in fact, be realized either through serial or parallel processes on a finer timescale. We therefore use the term *concurrent* processing in the present study to loosely denote processes that run together and to leave open the different possibilities of their fine-grained implementation. We develop a novel methodology to ask whether 1) we can find evidence for *concurrent* prospective action planning in human posterior parietal and premotor cortex by means of fMRI, and 2) if there is an individual capacity limit for concurrent planning due to the growing computational expense of multiple parallel computations, and how such a capacity limit could be quantified. With our new analytical approach, we develop task-specific optimal information-processing models with different constraints and regress the predicted information values with the recorded neuroimaging data to distinguish between serial and parallel motor planning during a delayed response task (DRT) that allows separating planning activity from sensory and motor processing [18].

In particular, the experimental paradigm of the present study was designed to contrast a serial memory-based planning strategy (null hypothesis H_0_: “delayed planning”) and a concurrent prospective planning strategy (hypothesis H_1_: “concurrent prospective planning”) to prepare for a goal-directed action consisting of a sequence of button presses during the delayed response phase (compare Figure 1). For example, an initial “cue” could indicate two potential target locations (e.g. either located in the left or right panel) so that during the ensuing delay period subjects could either prepare two possible action sequences for the upcoming response (H_1_) or simply remember the target locations and delay concrete action planning until the go-signal reveals the ultimate target location (H_0_). By modelling the task-specific information-processing efforts associated with each of the two respective strategies, we can then use a model-based regression to distinguish whether planning activity during the delay phase in any given area can be better explained by concurrent prospective planning (H_1_) or, alternatively, by delayed planning (H_0_).

**Figure 1:**
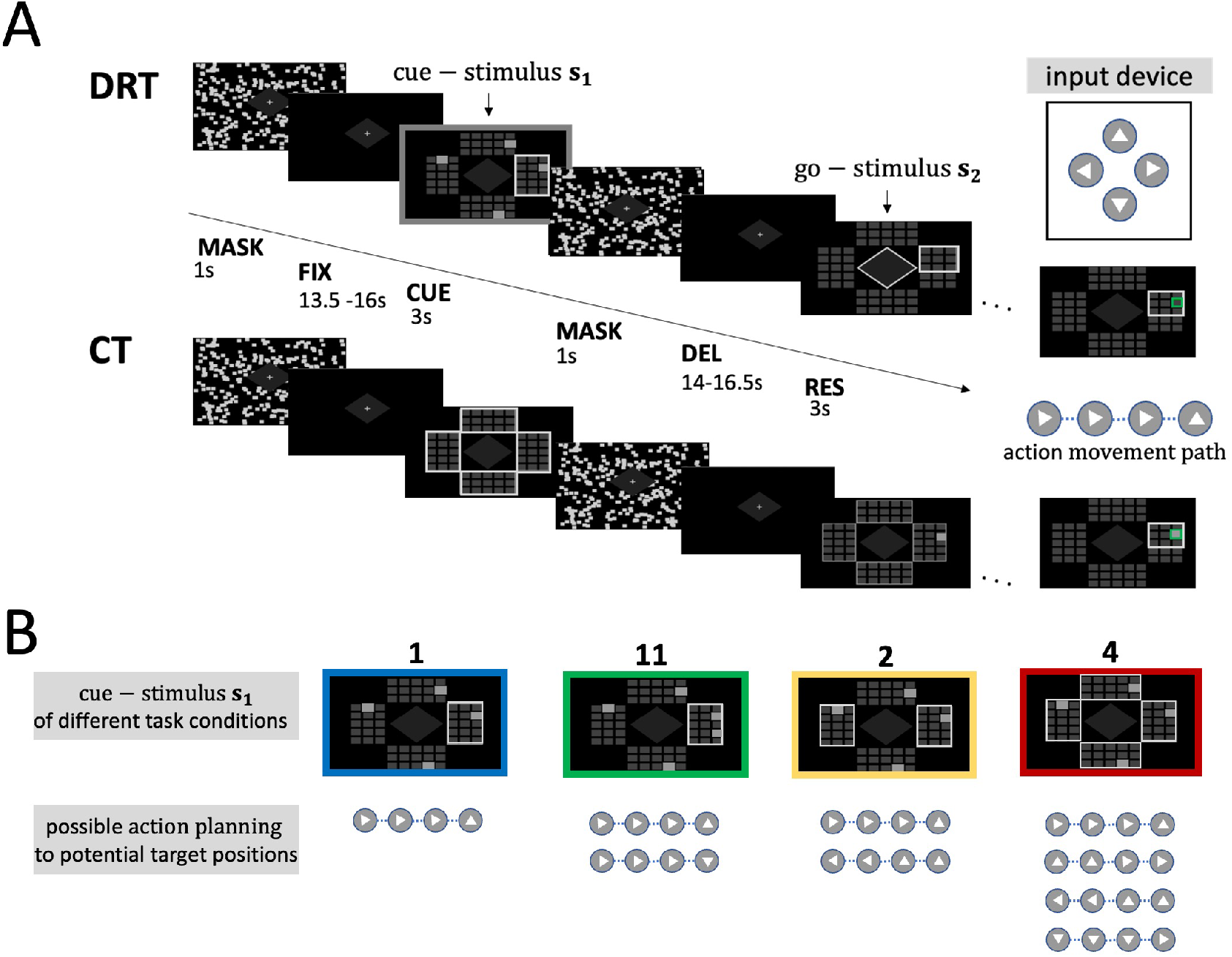
Experimental task design. (A) Trials of the motor planning conditions in the delayed response task (DRT) and control condition (CT) were randomized and had a similar timeline. Each trial started with a fixation period (FIX) of random duration, a cue phase (CUE), where a cue-stimulus *s*_1_ was presented, followed by a random dot pattern to mask afterimages of the CUE (MASK), a delay phase (DEL) of random duration for planning, and a response phase (RES). In the response phase (RES) the ultimate target location was revealed by a go-signal *s*_2_, namely a single grey frame around the relevant (part of the) respective response panel. Response trajectories were generated by subjects’ button presses on the illustrated input device. In the control task, no potential target cues were presented during the cue phase and therefore no goal-directed actions could be planned during the delay phase. In the response phase of the CT subjects were finally informed about the target location (single grey box). (B) The four target planning conditions differed in the number of potential target locations initially cued. Subjects could see possible target positions highlighted (grey boxes). Potential targets to consider for planning were indicated by a grey frame around the field areas encompassing the relevant possible targets. Subjects were instructed to plan a single movement sequence towards one target (‘1’), two partially overlapping sequences towards two potential targets (‘11’), two distinct movement sequences towards two potential targets in different panels (‘2’), or four distinct movement sequences towards four potential targets (‘4’) respectively and to execute the planned goal-directed movement as sequential button presses with the right thumb in the following phase, after the ultimate target was revealed by the go-signal. During the DEL phase, preparation for a goal-directed action could follow a serial memory-based planning strategy (null hypothesis H_0_: “delayed planning”), or a concurrent prospective planning strategy (hypothesis H_1_: “concurrent prospective planning”) where possible planning of actions as movement paths to each of the potential target locations could be anticipated.

Similar to previous studies that have quantified information-processing capacity in the context of working memory tasks [19, 20, 21, 22, 23, 24, 25, 26, 27], we use information theory in the model-based regression to quantify information-processing capacity in the context of motor planning. This can be achieved by designing optimal choice models with information constraints [28] that trade off the minimal amount of information processing required to achieve a certain goal, in our case planning to reach a target with a certain degree of accuracy. In previous studies [29, 30], we have investigated such optimality models with information bounds in purely behavioral motor planning tasks by manipulating both permissible reaction times and prior distributions over possible world states. In these experiments reaction time was used as a proxy for information cost underlying any process of uncertainty reduction. Here we apply such optimal information-processing models to human fMRI activity in order to reveal whether brain areas engaged in the planning of goal-directed actions resort to a delayed planning strategy (H_0_) or, alternatively, to concurrent prospective planning (H_1_). In addition we used these models to estimate planning-related capacity limits. Based on this approach we show that brain activity in a parietal-frontal planning system and the cerebellum is consistent with bounded optimal information flow implementing concurrent prospective action planning.

## 2 Results

### Behavioral performance analysis

In our delayed-response-task (DRT), 19 subjects (11 females, 8 males, average age = 27.5) were shown an initial cue stimulus with potential target locations that were highlighted in four concentrically ordered response panels. They were verbally instructed to plan movement sequences to potential targets during the subsequent delay phase, and to execute one of these sequences, when shown the go-signal as a selection stimulus during the response phase (see Figure 1 B). Movement sequences could vary in sequence length (2-step, 3-step and 4-step condition) and consisted of up/down/left/right button clicks to move a cursor on a screen. To enforce planning during the delay phase, the go-signal did not indicate the actual selected target directly, but only indicated the relevant (part of the) response panel. In other words, subjects could uniquely identify the selected target only if they remembered the information provided by the initial cue stimulus. In total, our task design contrasted 3×4 DRT conditions, that did not only vary both in the movement sequence length (2, 3, or 4), but also in the number of potential targets (1, 2 or 4 targets in distinct response panels or condition ‘11’ with 2 targets in the same panel). Consequently, the 12 DRT conditions were distinguished by different information costs due to multiplicity of potential panels, potential targets and sequence lengths.

We assessed subjects’ behavioral performance in terms of reaction times, movement times and error rates and compared to a control condition (CT), where the initial cue stimulus did not contain any information about the target location but the relevant target location was explicitly cued during the response phase. Therefore subjects did not need to memorize or plan during the delay phase of the control condition.

We found that reaction times increased with the number of potential targets (*p* < 0.0001, 2-way rmANOVA with factors condition [‘1’,‘11’,‘2’,‘4’,‘CT’] and sequence length: linear effect of condition – see Methods section for details on the statistical analysis), suggesting that subjects engaged in more complex planning as the number of potential targets increased (see Figure 2 A). In contrast, sequence length had no significant effect on reaction times (p = 0.441).

**Figure 2:**
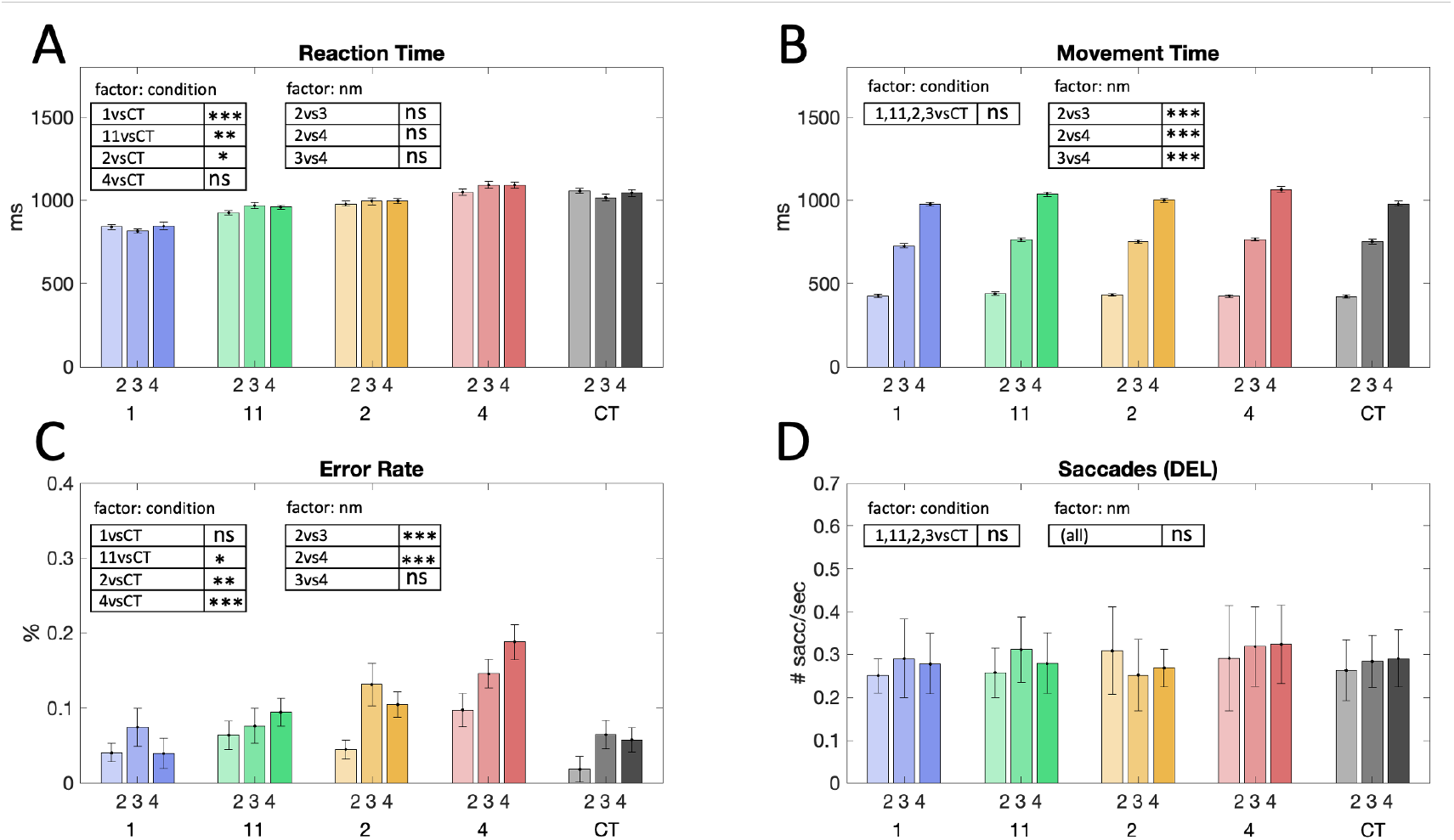
Behavioral performance. Results are reported as the mean over subjects’ individual task performance and normalized standard error (according to [31] to eliminate the between-subject variance, which does not contribute to the within-subject effect of the study). Results of the one-sided pairwise post-hoc comparisons of the respective marginal means are indicated with *** for *p* <= 0.001, ** for 0.001 < *p* <= 0.01, and ns for non-significant results (*p* > 0.05). (A) Reaction times (RT) and (B) movement times (MT) were calculated as averages across individual subjects’ means ± SEM. Reaction times were significantly decreased for easy planning conditions (‘1’, ‘11’ and ‘2’) compared to the control condition CT, which indicates a benefit due to planning (‘1’ vs. CT, *p* < 0.0001; ‘11’ vs. CT, p = 0.0002; ‘2’ vs. CT, *p* = 0.0167). RT were higher in the more complex planning condition (‘4’) and did not significantly differ compared to CT (‘4’ vs. CT, *p* = 0.933). Differences for varying sequence length were found for MT but not for RT. (C) Error rates averaged across subjects indicated varying task difficulty over conditions revealed by an increase of target misses for conditions with higher task uncertainty (*p* = 0.024 for 11 vs CT; *p* = 0.005 for 2 vs CT; *p* < 0.001 for 4 vs CT) and between 2- and 3-step conditions and 2- and 4-step conditions (*p* < 0.0001 for 2 vs 3; *p* < 0.0001 for 2 vs 4). (D) During the delay phase there was no significant difference between the average number of eye saccades and therefore saccades cannot explain benefit in reaction times or fMRI activity in planning-related areas.

Compared to the DRT conditions, the control condition CT did not allow for any planning during delay. In case of concurrent prospective planning (but not for delayed planning), we therefore expected to reveal reaction time benefits in DRT as compared to CT. Indeed, a priori one-sided pairwise t-tests of reaction times averaged over the sequence length reveal behavioral benefits in planning conditions compared to CT. Reaction times in the less costly planning conditions ‘1’ ‘11’ and ‘2’ were significantly reduced as compared to CT (‘1’ vs.CT, *p* < 0.0001; ‘11’vs.CT, *p* = 0.0002; ‘2’ vs. CT, *p* = 0.0167, one-sided pairwise t-test). For the more complex planning conditions ‘4’ no such reaction time advantage was found (‘4’vs. CT, *p* = 0.933, one-sided pairwise t-test) (see Figure 2 A). These results also hold up when conducted as a post hoc test with a Bonferroni-Holm correction for multiple comparisons (4 comparisons with a family-wise error rate of *α* ≤ 0.05), where again conditions ‘1’ ‘11’ and ‘2’ show a significant reaction time benefit (‘1’vs.CT, *p* < 0.0125; ‘11’vs.CT, *p* < 0.0167; ‘2’ vs. CT, *p* < 0.025), but not condition ‘4’ (‘4’ vs. CT, p > 0.05).

As to be expected, there was a significant effect of sequence lengths on movement times as shown in Figure 2 B (*p* < 0.0001, 2-way rmANOVA with factors condition [‘1’,‘11’,‘2’,‘4’,‘CT’] and sequence length: linear effect of sequence length). There was no linear effect of condition on movement time (*p* = 0.449).

Subjects’ error rates are shown in Figure 2C. There was a significant influence of the factor condition [‘1’,‘11’,‘2’,‘4’,‘CT’] (*p* < 0.0001, 2-way rmANOVA) and sequence length (*p* = 0.018). In the CT, the target was explicitly shown after the delay, which perhaps explains the lowest error rates in this condition. In contrast, the DRT conditions required memorization of the cue stimulus, which allowed for additional errors and thus led to higher error rates in the more complex DRT conditions.

Finally, during the delay phase the number of saccades per second did not significantly differ between the different planning conditions and control conditions (see Figure 2 D). This absence of difference in eye movements during the delay phase will be important in the next section, where we study fMRI activity during the planning delay period and need to exclude additional effects on fMRI from eye movements.

### fMRI analysis

Brain activity during task performance was measured by means of fMRI in a 3T MRI scanner. We fitted a general linear model (GLM) to the activity time series data measured in each subject and in each voxel of the brain. In this GLM we modeled each of our 3×4 conditions and separately for cue, delay and response phase (for details see methods). We used the resulting regression models to determine regions of interest that show significant functional delay-related activity on the group level. To this end, the delay-related signal estimates of all 12 DRT conditions were contrasted against the control condition CT in each individual subject. The resulting group map is depicted in Figure 3 C. Our second-level group analysis showed significant differences across subjects in the planning-related areas of the left and right (l/r) posterior parietal and premotor cortex that we hypothesize to be engaged in prospective parallel planning, namely SPLl, SPLr, antIPSl, antIPSr, PMdl, PMdr. In addition, delay activity was exhibited in dorsolateral prefrontal cortex (DLPFCr), the anterior insular cortex (AICl, AICr), supplementary motor area (SMA), as well as the lobules 6 and 8 of the cerebellar hemispheres (cer6l, cer6r, and cer8r). While we centered our analyses on the earlier areas in posterior parietal and premotor cortex as they have been previously shown to contribute to concurrent prospective planning in monkeys (compare introduction), we also performed explanatory analyses of all other aforementioned areas. For further details on how these regions of interest (ROIs) were selected in individuals, please see the Method section. For each of our predefined ROI areas, we extracted GLM parameter estimates reflecting delay-related changes in fMRI BOLD signal amplitudes in different task conditions of varying planning complexity. In the following we chiefly focus on area SPLl, as an exemplary target ROI. For results of different ROIs – left cerebro-cortical ROIs contralateral to the effector and ipsilateral cerebellar ROIs in the right hemisphere – please refer to Supplementary Figure S1. In area SPLl, delay-related GLM parameter estimates differed significantly between individual planning conditions, distinguishing both the number of potential targets (p < 0.0001, 2-way rmANOVA with factors condition and sequence length) and sequence length (p = 0.001) but there was no significant interaction of the two factors. The relative changes of the fMRI signal over time due to planning during the delay phase compared to a pre-stimulus baseline can be seen for the different conditions in Figure 3 A. As expected, the activity in SPLl was lowest during the control condition CT that does not provide any information for planning. The activity increased with the complexity of the planning problem with the highest activity for the target condition ‘4’ with sequence length 4. As a control ROI, primary visual cortex V1l – that is supposedly not involved with motor planning – showed no such modulation (see Figure 3 B). Note that due to the slow BOLD haemodynamic response function (HRF), visual processes and eye movement related activity during cue presentation could influence the time courses of fMRI-activity in the early delay phase of planning. Differences in pure planning activity for different task conditions in isolation are thus only reflected by the time courses in the late delay phase. Hence, at first glance, planning related activity in SPL and in all other planning-related ROIs (compare Supplementary Figure S1 with PMdl, antIPSl, DLPFCl, AICl, cer6r, cer8r, SMA) seemed compatible with concurrent prospective planning.

**Figure 3:**
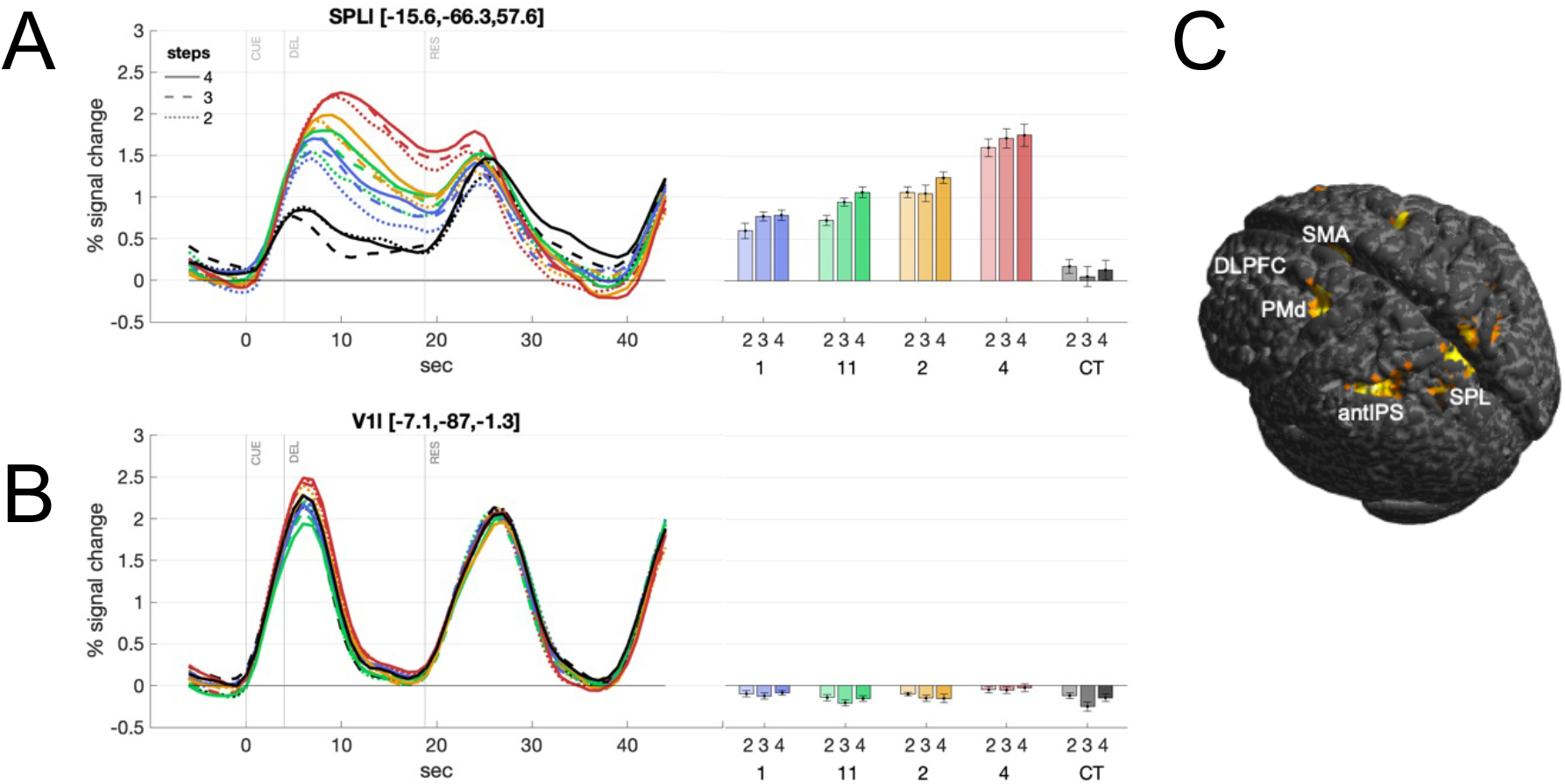
fMRI activity related to motor planning. (A) Time course of fMRI activity as percentage of signal change normalized to pre-stimulus baseline and delay-activity estimates (beta parameter from GLM) in each planning condition in planning related area SPLl show increasing planning-related BOLD amplitudes with increasing task complexity. We report across-subjects averages and within-subjects variance as the normalized standard error (according to [31]). Time course activity and GLM estimates of further ROIs – contralateral cortical ROIs (left) and ipsilateral cerebellar ROIs (right) – are provided in Supplementary Figure S1. Average MNI-coordinates (x, y, z in mm) are provided for each ROI. Statistical results are indicated with *** for *p* <= 0.001, ** for 0.001 < *p* <= 0.01, * for 0.01 < *p* < 0.05, and ns for non-significant results (*p* > 0.05). (B) In a ‘non-planning ROI’ (V1l) BOLD signal changes were not significantly different between conditions. (C) Second-level activation map shows significant delay-related fMRI activity across subjects in planning related areas (see Results for details) when contrasting DRT conditions vs. the control condition CT (*p* ≥ 0.05 family-wise error (FWE)-corrected for multiple comparisons).

### Model analysis

In our delayed-response task, information about the ultimate target location (the hidden world state 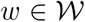) is revealed successively by the two stimuli *s*_1_ and *s*_2_: the initial cue stimulus 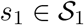 indicates all potential target locations; the second go-stimulus *s*_2_ resolves the remaining uncertainty about the actual hidden target *w* (see Figure 4). Accordingly, the overall process can be described with two information quantities (*I*_1_ and *I*_2_), that are dependent on the respective task condition (e.g. the number of potential targets and the length of the action sequence that leads to these targets). In a first step, the information *I*_1_ is needed to form and maintain a memory 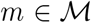 of the initial cue, which is needed to infer the world state when seeing *s*_2_ after the delay. The process of memory formation therefore partially resolves some uncertainty about *w* given the amount of information that *s*_1_ provides. This process and its associated information *I*_1_ is identical in both planning strategies, delayed planning (**H**_0_) and concurrent prospective planning (H_1_).

**Figure 4:**
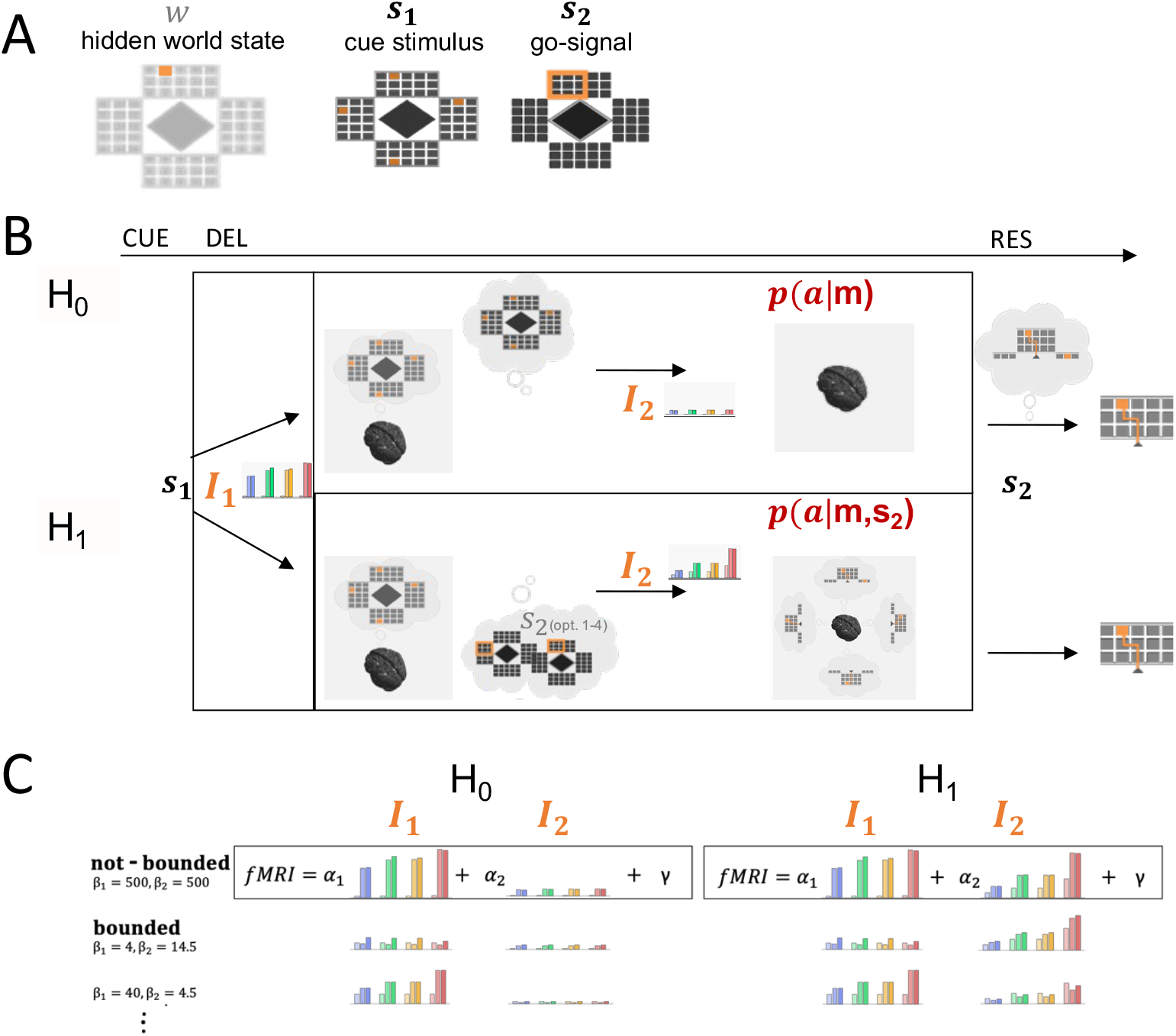
Information theoretic model of planning with information constraints. (A) Information about the hidden world state 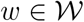 (the exact target location) is revealed in two steps during the experimental trial. The initial cue stimulus 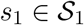 indicates potential target locations while the later go-signal *s*_2_ resolves any remaining uncertainty about the actual hidden target *w*. (B) When seeing *s*_1_, the decision-maker can in a first step form a memory 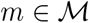 which is required to infer the world state when seeing *s*_2_ after the delay and, therefore, is then able to appropriately plan and finally select an action 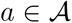 that corresponds to a movement path representation. Both hypothesis (H_0_ and H_1_) contrast the predicted information *I*_1_ and *I*_2_ in all 12 DRT conditions for a delayed planning strategy and concurrent prospective planning strategy during the delay (DEL). H_0_: During the delay phase only memory *m* is used to reduce uncertainty about actions. H_1_: Concurrent prospective planning of actions during delay phase requires the anticipation of all possible selection stimuli *s*_2_ and planning movement sequences concurrently. *I*_2_ therefore is higher in H_1_ than in H_0_ and, moreover, varies more strongly with movement sequence length. (C) Model Comparison. We regressed theoretical memory and prospective planning information with measured fMRI activity in relevant brain areas during the delay phase and tested two hypotheses: H_0_, where information-processing merely requires uncertainty reduction based on memory formation *m* and H_1_ with higher information-processing effort required for prospective planning to anticipate possible future movements. Theoretical information values *I*_1_ and *I*_2_ for different memory and planning capacities, defined by model parameters *β*_1_ and *β*_2_, were regressed (using regression coefficients *α*_1_ and *α*_2_) with the measured fMRI BOLD activity. For lower *β*_1_ and *β*_2_, memory and planning capacity accordingly is more limited. We regressed the information values for different degrees of boundedness and compared the best model fits with the maximal information values in the case without bounds, where *β*_1_ and *β*_2_ were chosen maximally (*β*_1_ = 500, *β*_2_ = 500).

In the second processing step, uncertainty about actions gets reduced. This is quantified by additional information *I*_2_, that, however, differs for the delayed planning and the concurrent prospective planning strategy. In the delayed planning scenario (H_0_), only memory *m* is used to reduce uncertainty about actions and further information processing is suspended until the go-stimulus *s*_2_ gives the final information indicating the hidden world state. For concurrent prospective planning (H_1_), however, all potential target locations are considered and information costs are added up for planning all respective action sequences towards these targets. *I*_2_ therefore is generally much higher for concurrent planning and is also increasing more strongly with the number of potential targets and movement sequence length for a concurrent prospective planning strategy as compared to a delayed planning strategy (see Figure 4C). This difference in integrated information costs *I*_1_ and *I*_2_ allows us to probe whether fMRI activity in the delay period does reflect concurrent prospective planning (hypothesis H_1_) or serial delayed planning (hypothesis H_0_). Moreover, while prospective concurrent planning brings essential benefits by enabling subjects to react more quickly to an upcoming scenario [32], it is immediately clear that this type of concurrent planning cannot be scaled up indefinitely and that there must be bounds on the amount of information processing. For this reason we also assess these capacity limits of information processing.

In order to formally compare the two planning hypotheses – concurrent prospective planning (H_1_) against the null hypothesis of delayed planning with uncertainty reduction only based on past sensory information (H_0_) – we compared the fMRI activity modulation during the delay phase over different experimental conditions to the expected information modulation determined for both hypotheses from an optimal decision-making model with information constraints (see Figure 4). For a detailed description of both hypothesis please see the Methods section. Assuming motor planning during the delay phase only in terms of uncertainty reduction based on memorized information of the perceived cue stimulus, the decision-maker requires information cost *I*_1_ to first update the internal memory. Correspondingly the action uncertainty gets reduced from a prior over all actions to a posterior over potential actions given the memory, which requires an additional information cost *I*_2_. We contrasted this null hypothesis H_0_ with the assumption of concurrent prospective planning, where separate action plans are made for all potential targets. In this alternative hypothesis H_1_, the memory update is identical to H_0_ (same information cost *I*_1_), but the information cost required for the reduction of uncertainty over actions is now composed of the sum of information costs to form each possible motor plan (*I*_2_). The exact amounts of information (*I*_1_ and *I*_2_) depend on the capacity of the information channels defined by model parameters *β*_1_ and *β*_2_ for the memory and the action process. Three example profiles of information modulation with different capacities are shown in Figure 4 C.

### Model comparison

To find out which hypothesis explains our data best, we established (non-negative) multilinear regressions between the fMRI activity modulation during the delay period and the two information modulations predicted by each of the two hypotheses. For both hypotheses, we first calculated the expected information costs for possible settings of information capacities and we regressed these theoretical information modulations with the respective subject-specific GLM activity estimates over the 12 DRT conditions for all ROIs separately. Figure 5 A shows, exemplarily for brain area SPLl, the *R*^2^-values of the regressions for the best fitting information capacities for each subject under the two hypotheses (which we refer to as “bounded” in the Figures 4 and 5). A within-subject comparison of the *R*^2^-values for the two hypotheses showed that the information profiles under the prospective planning hypothesis H_1_ provide a significantly better explanation of the fMRI modulation than the delayed planning hypothesis H_0_ in SPLl (SPLl: *p* < 0.0001 rmANOVA) as well as in all other planning related ROIs, but not in control ROI V1l (compare Supplementary Figure S2 and Supplementary Table 2). The superiority of hypothesis H_1_ can also be confirmed by leave-one out cross-validation, as it allows for significantly better predictions of fMRI activity in all relevant brain areas, except V1l and M1l (compare Supplementary Table S10). For further details on the statistical analysis see the Methods section. Moreover, we applied a nested model F-test statistic to both hypotheses to find out whether the predicted planning-specific information *I*_2_ contributed significantly to the multilinear regression of the fMRI signal. We found that for most of the subjects (16 from 19 for SPLl) under the prospective planning hypothesis H_1_ the planning information *I*_2_ improved the regression of the signal modulation significantly (see Figure 5). In contrast, the nested model tests for H**0** revealed that model regressions did not improve significantly by taking *I*_2_ into account.

**Figure 5:**
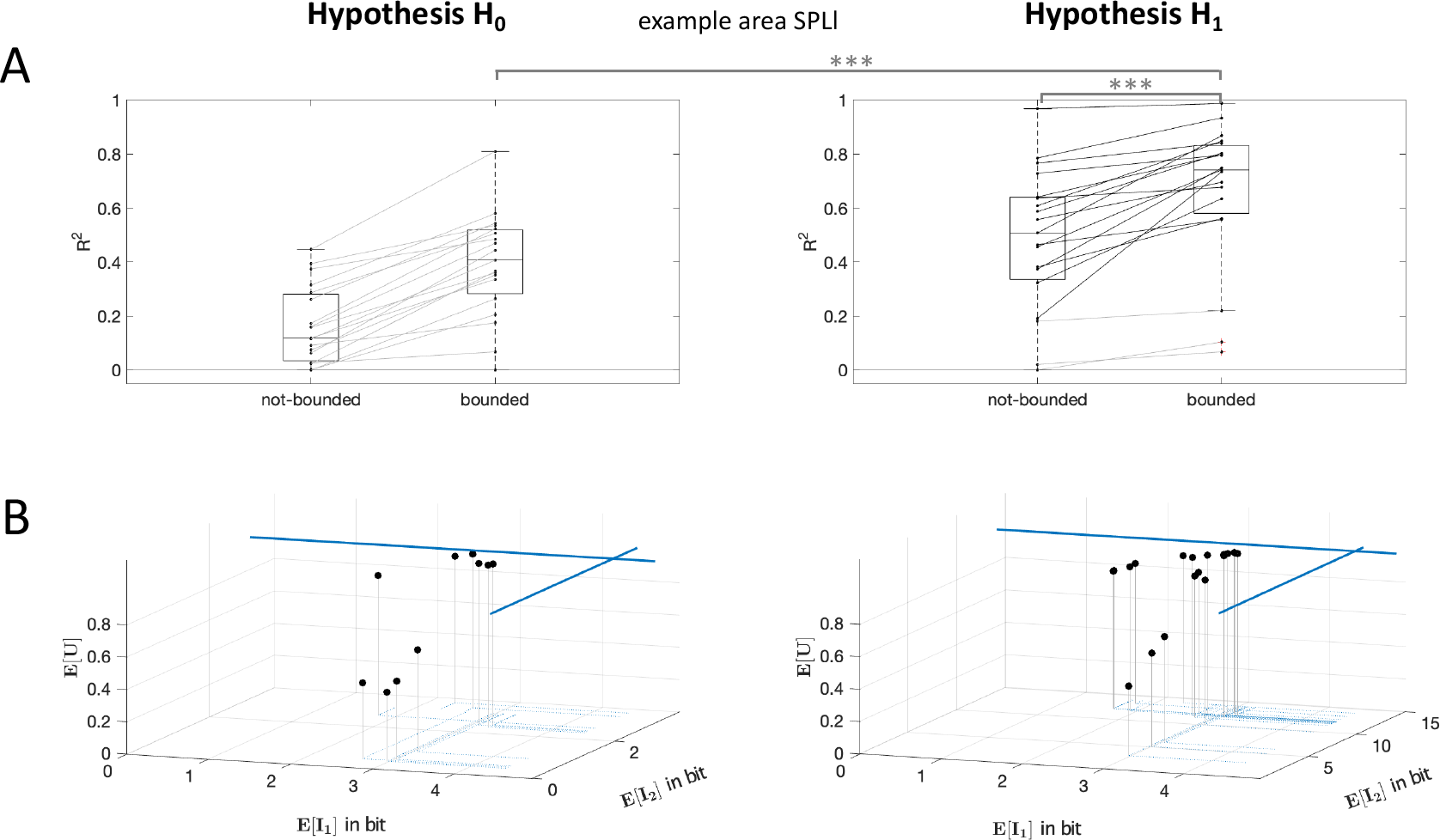
H_0_ and H_1_ model comparison. (A) *R*^2^-values of the regressions for the best fitting information capacities for individual subjects for H_0_ and H_1_ are compared to not-bounded models. Statistical results of rmANOVAs are indicated with *** for *p* <= 0.001. (B) Resulting theoretical quantitative predictions of the expected information costs and expected utility over all task conditions are plotted for individual subjects. Expected values of information costs *E*[*I*_1_] and *E*[*I*_2_] of subjects’ bounded model fits to the fMRI correlates was lower than for not-bounded model (blue lines). This gives hint to prospective planning activity with subject-individual planning capacities.

To address the question in how far concurrent planning is limited in its capacity, we compared the *R*^2^-values of the regression with an optimal decision-making model without capacity bounds that we obtain as a limit case of maximum capacity (which we refer to as not-bounded in Figures 4 and 5). We found that the proportion of explained variance of measured brain activities in SPLl as well as in all other planning related ROIs was significantly increase for model predictions with subject-individual bounds (SPLl: *p* < 0.0001 rmANOVA, compare Supplementary Figure S2 for all other ROIs) compared to an unbounded maximum capacity model. This suggests that judging from the modulation of the fMRI signal alone, it seems that subjects were limited in their information processing. As a control, the *R*^2^-value of the information modulation in primary visual area V1l was close to zero (see Supplementary Figure S2). Note that our within-subject comparisons of the *R*^2^-values for the two hypotheses H_0_ and H_1_ remain stable also for assuming a non-linear relation (quadratic, sigmoidal, logarithmic) between fMRI correlates and predicted information values. Regressions produced similar results (see Supplementary Figure S4 and Supplementary Table 1 for p-values of rmANOVAs). Similarly, under the prospective planning hypothesis H_1_, the planning information *I*_2_ improved the regression of the signal modulation significantly (nested model F-test statistic significant for 15 of 19 subjects for SPLl) and *R*^2^ values regressing model predictions with subject-individual bounds were significantly higher compared to an unbounded maximum capacity model. Again this result can be confirmed by leave-one out cross-validation comparing prediction errors of fMRI activity with and without information bounds (compare Supplementary Table S10).

The individually fitted information bounds under the two hypotheses H_0_ and H_1_ can be seen in the 3d graphs shown in Figure 5 B. The expected information costs over all conditions *E*[*I*_1_] and *E*[*I*_2_] of subject-individual best model fits were lower compared to the predicted information boundaries *E*[*I*_1_] an *E*[*I*_2_] when assuming no bounded information capacities for *I*_1_ and *I*_2_ (blue lines). Our comparison of the expected information values for the bounded and not-bounded model indicated, that for all subjects mainly information capacity for prospective planning *I*_2_ was reduced whereas information for memory *I*_1_ was reduced but still closer to the predictions of the model with the not-bounded assumption. Expected information values for subject-specific model fits to activity in all other ROIs can be found in the Supplementary Material (Table 3-Table 6). Assuming a simple 0/1-utility function *U* for our task, that assigns a utility value of 1 to actions *a* that reflect a target hit the value 0 otherwise (see Methods section), the expected utility *E*[*U*], shown on the z-axis, was individually different for the decision-making models (see Supplementary Tables 7 and 8) and is higher for subjects with higher capacity limits for *I*_1_ and *I*_2_.

Figure 6 A shows a comparison of *R*^2^-values as the average proportion of explained variance of measured fMRI signals in different ROIs resulting from the regressions for the best fitting information capacities for individual subjects for H_1_. For each individual ROI, we compared *R*^2^-values using one-sided non-parametric permutation statistics. For all planning related areas we found *R*^2^-values to be significantly lower than for control areas M1l and V1l (*p* < 0.0001). Further, only for fMRI modulations in SPll, the model fits could provide significantly better explanations compared to antIPSl (*p* = 0.0013), PMdl (*p* = 0.0172), DLPFCl (*p* = 0.0036), cer8r (*p* = 0.0094), and AICl (*p* = 0.0051).

**Figure 6:**
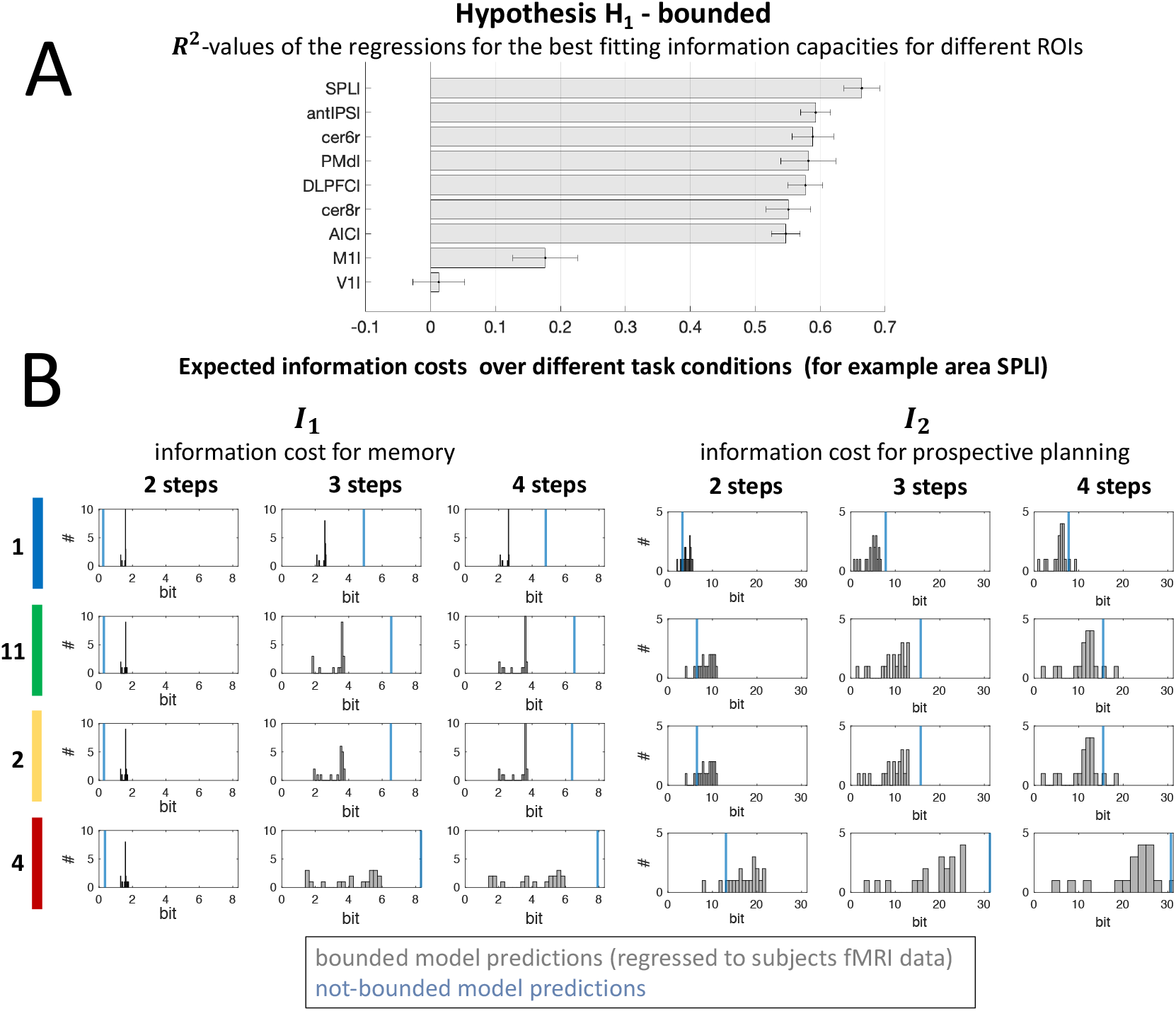
Bounded prospective planning (H_1_) regression. (A) ROI comparison ranked by *R*^2^-values of the regressions for the best fitting information capacities for individual subjects for H_1_. Results are reported as the mean over *R*^2^-values for subjects’ individual best model fit for H_1_ and normalized standard error (according to [31] to eliminate the between-subject variance, which does not contribute to the within-subject effect of the study). (B) Theoretical expected information costs *I*_1_ (left) and *I*_2_ (right) of prospective planning is based on model parameters determining memory and planning capacities fitted for all individual subjects (hypothesis H_1_ “bounded”). Histograms represent the frequency distributions of the information costs, dependent on task conditions. Information values varied between bounded model predictions regressed to subjects fMRI data compared to the model predictions in the not-bounded case (in blue). Note that generally subjects lie below the information cost of the not-bounded decision-maker and only for the most simple 2-step conditions subjects’ information costs for memory (*I*_1_) deviate in the different direction from the predictions in the not-bounded case. This results from the specific task design and the fact, that bounds are provided on the total expected information. Decision-makers with not-bounded planning capacities can optimally assign resources to more difficult conditions and save memory costs for 2-step conditions, because no uncertainty about the target location will remain when *s*_2_ is revealed given that all actions are planned prospectively under a high information resource for *I*_2_.

To investigate the impact of limited information processing for concurrent prospective planning in individual task conditions with different complexity (i.e. number of potential targets and sequence length), we illustrated frequencies of information costs *I*_1_ and *I*_2_ for subjects’ best model fits in a histogram representation for each condition (see Figure 6B; for comparison of H_0_ hypothesis please refer to Supplementary Figure S3). We contrasted the frequency distributions with the information predictions for a perfect decision-maker without information processing bounds (blue line). Deviations from the information with maximal capacities show evidence for bounded planning. It can be seen that subjects generally lie below the information bound of the perfect decision-maker.

Finally, we test the models’ predictive power for behavioral performance from fits to fMRI activity during the planning phase. Naturally, one may not expect a perfect prediction, since ultimate behavioral performance depends on many other factors, both detrimental ones like motor execution noise and imperfect planning as well as beneficial ones like additional planning following the final cue. To obtain a prediction for subjects’ behavior from individual fMRI activity, we compute the predicted expected utilities according to equation (4) based on the probabilities we obtained from the optimal information-fMRI fit in Figure 5 under both hypothesis H_0_ and H_1_. These predicted planning performances were then compared to subjects’ actual behavioral performances across the experiment in terms of error rates (see Figure 7). Correlation of the experimental utilities (1 – error rate) and the theoretical *E*[*U*] values were analysed by linear regression. Only for the concurrent planning hypothesis H_1_, we found a significant correlation (*p* = 0.0238) between measured data and predicted outcome of the model. For the null hypothesis H_0_, subjects’ expected utility estimates were not significantly correlated with the measured behavioral performance. Thus, we may conclude that our optimal information model has significant predictive power with respect to subjects’ behavioral performance from fMRI activity recorded during the planning phase in our delayed-response task.

**Figure 7:**
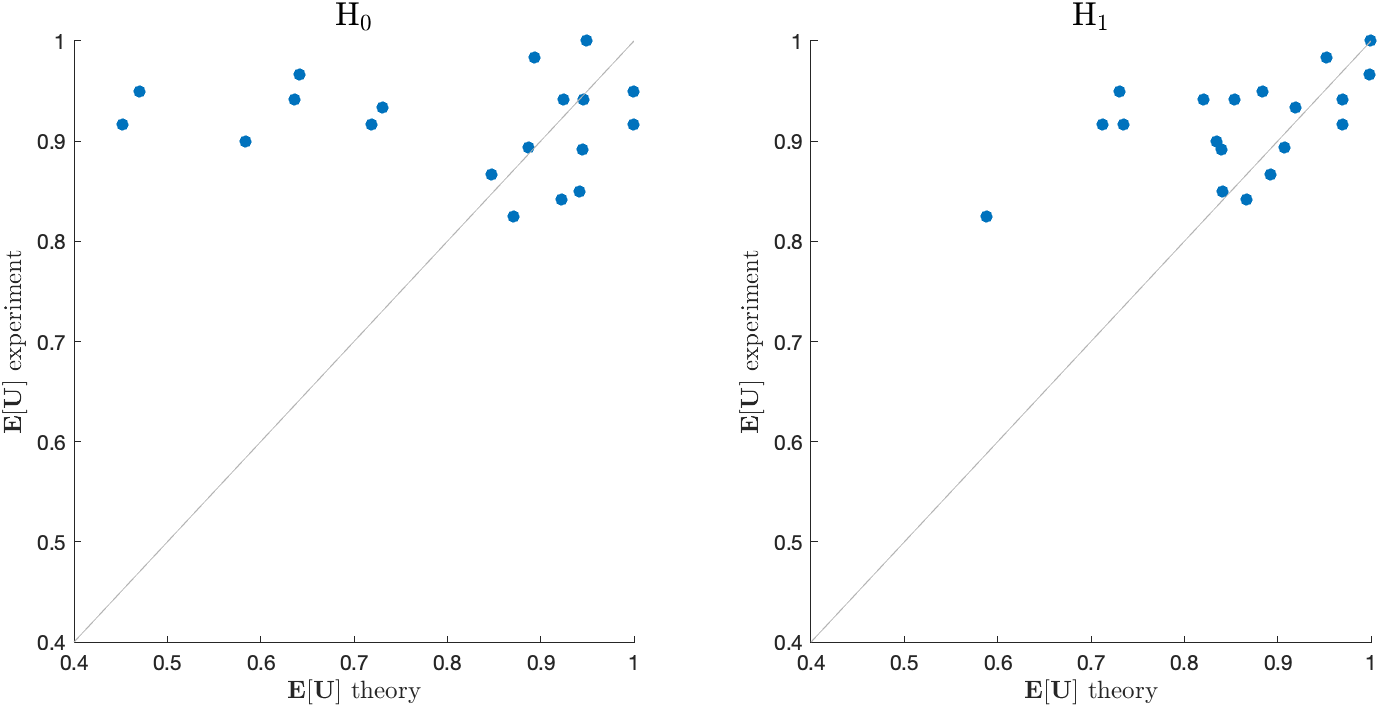
Comparison of experimental and theoretical performance. Linear regression analysis of theoretical *E*[*U*] averaged across model predictions over all planning related ROIs (DLPFC, PMd, SPLl, antIPS, AIC, cer6, cer8; compare Figure 6A) under both model hypothesis H_0_ (left) and H_1_ (right) and experimental utility (1 - error rate) for the group of subjects. Whereas under model hypothesis H_0_, individual *E*[*U*] resulting from model fits and subjects fMRI data was not significantly correlated to measured experimental utility (linear regression with *p* = 0.555, correlation coefficient *ρ* = −0.145 and regression slope *m* = −0.535), under H_1_ a significant correlation between measured data and estimated data was found (*p* = 0.0238) with correlation coefficient *ρ* = 0.516 and regression slope *m* = 1.171. Better experimental performance in individual subjects than theoretically optimal performance can be plausibly explained by the fact that after the delay phase subjects could still improve their behavior through additional planning following the second cue, whereas the theoretical optimum is computed for planning only with the first cue. Linear regression with shuffled experimental data was repeated for result validation. For shuffled data, only 3.5% of correlations were significant with correlation coefficients *ρ* in the range of [−0.671, 0.608] and regression slopes *m* in the range of [−1.524, 1.380].

## 3 Discussion

Using fMRI and information-theoretic modeling, we have investigated neural correlates of information processing costs during concurrent motor planning in the human motor system [18]. We manipulated informational complexity of movement planning by varying both the temporal extension of the movement sequence length as well as the degree of concurrency (the number of potential targets). We compared how much of the variance of the fMRI signal in specific brain areas (i.a. in PPC and PMd) could be explained when assuming concurrent prospective planning for multiple potential targets (hypothesis H_1_) and when assuming no prospective planning, but only uncertainty reduction based on past sensory information (hypothesis H_0_). We found that the theoretical information costs of concurrent prospective planning explained fMRI profiles across conditions significantly better than assuming a memory-based strategy (hypothesis H_0_) that delays all anticipatory planning activity into the future until a second sensory stimuli (the go-signal) resolves the remaining uncertainty. Finally, we found that the bounded optimal planning performance error predicted for individual subjects by our concurrent model from fMRI data, correlated significantly with subjects’ actual performance error during movement execution. Our results therefore add to a growing body of evidence that suggests the use of concurrent prospective planning in the parietal-frontal planning system when faced with multiple potential futures [33, 34, 7, 8, 35, 36, 10, 2, 37].

While the concurrent planning of multiple potential futures comes with a reaction time advantage whose behavioral consequences could be positively selected by evolution, it is also immediately clear that it cannot scale to scenarios with arbitrarily large amounts of uncertainty due to an excessive computational burden. Therefore, one would naturally expect a trade-off between the degree of concurrency in motor planning and the amount of information processing required. Evidence for such a trade-off has been previously reported in behavioral studies [38], where it has been shown that competition between movement plans increases motor variability, thus suggesting a shared resource for movement planning. In our experiment, this could be seen both in the behavior and the fMRI BOLD signals. Behaviorally, we noted an increased error rate for conditions with high degree of concurrency, suggesting a limitation in precision for concurrent planning. Neurally, we found that the best model correlates are characterized both by the assumptions of concurrent planning and limited information capacity. In particular, we found in our nested model analysis for the two information profiles regarding memory and planning that adding the planning information significantly improved the variance explained in the fMRI signal.

### Serial vs. parallel processing

In our study the null hypothesis H_0_ has been aligned with more traditional theories of sensorimotor processing whose point of view is that perceptual, cognitive and motor processes take place successively and only a single motor program is prepared in the end [5, 6]. It should be noted that recent experimental evidence supports this view regarding both behavioral and neural data in monkeys and humans performing reaching movements towards spatial targets [12, 13]. However,their experimental paradigm differs from ours where motor planning relates to target-directed cursor movements controlled by sequential button presses.

Neural correlates of planning in MEG for sequential finger movements (yet without spatial targets) have previously been studied for example in the context of “competitive queuing” [39] and found evidence for parallel representations of sequence positions. Our study results add to these findings, in that it demonstrates neural correlates of multi-target planning in a target-directed movement sequence task in humans. Our model approach thereby can be seen as first evidence for concurrent prospective planning of sequential target-directed actions in human posterior parietal and premotor cortex based on an optimal information processing model that allows to distinguish between different hypotheses of serial and concurrent planning. Our results suggest that multiple plans are formed during the delay phase, as fMRI activity correlates with planning costs summed over potential actions. However, ultimately it remains open whether these plans are formed simultaneously or whether they are rehearsed one-by-one on a finer timescale. Moreover, it might be difficult to generalize our results to determine whether parallel plans are formed in more natural contexts. Whether or not there is a representation of alternative action plans might ultimately depend on individual task requirements [40], and – as we show in our study – on individual information capacities.

Previous neural studies that have favored the hypothesis of parallel motor planning have been mostly conducted in monkeys. Cisek and colleagues [34] have found, for example, that PMd activity in a two-target condition reflects presence and relative location of both targets, and that once the right target is indicated by a nonspatial cue, the corresponding directional signal is increased, while the other one is suppressed, thus, suggesting two concurrent motor plans vying for execution (but see Dekleva et al. 2018 [13] for an alternative interpretation). Similarly, Klaes and colleagues [10] have found that neurons in the frontoparietal reach areas of monkeys simultaneously represent two different movements associated with two different sensor-action rules, when it was made unclear to the monkey which of the rules to apply. In a similar vein, Cui and colleagues [2] have found in a nonspatial effector choice task (saccade vs. reach) that the parietal reach region encodes both potential action plans, whereas the dorsal area 5d only reflected the ultimately selected effector. A recent fMRI study [37] also reported supportive evidence for the affordance competition hypothesis in humans, demonstrating that task-relevant information biases activity of primary motor regions before movement onset and that coupling with fronto-parietal regions increased when the evidence for competing actions was similar. Other studies that have supported the affordance competition hypothesis in humans have mostly concentrated on behavioral features suggestive of concurrent planning [41, 15].

Other studies [1, 42, 43] have argued that decisions between different options are made upstream in prefrontal brain areas and are independent of the particular sensori-motor contingency of the task, arguing against the notion of concurrent action planning. However, decisions in these tasks usually involve choices between goods (e.g. apple vs. banana), where the effector is not directly relevant for the choice, even though different action costs can be integrated within the valuation of different options [42]. This is in contrast to action-based decisions like in our study, where the choice is tightly linked with the effector through the spatial nature of the stimuli. Our study clearly falls in the latter category and fits with previous results compatible with concurrent planning. In how far these results can be generalized to decisions between nonspatial stimuli with different values is an open question that might require the careful comparison of different stimulus and task designs.

In contrast to neurophysiological studies in monkeys that showed the representation of multiple action plans in PCC and PMd [34, 10], which is compatible with our results, we additionally find neural correlates of multi-target planning costs also in other brain areas such as the cerebellum and the AIC. A contribution of these areas in our task is not unexpected. First, cerebellar activation related to motor control and movement sequence processing [44] is mainly found in intermediate and lateral lobules VI and VIII [45, 46], which is consistent with the localization of our cerebellar ROIs. More importantly, recent electrophysiological evidence from monkeys performing a delayed-response task engaging hand movements clearly revealed preparatory activity in the ipsilateral hemisphere of lobules V and VI, which markedly mirrored activity profiles found in premotor cortex [47]. Note that one reason why previous fmri research using comparable tasks did not report planning activity in the cerebellum is that this structure was simply not included in the scanning volume (compare Lindner et al. 2010 [18]). Second, the AIC is also active during delayed response tasks and for a variety of task conditions with and without motor planning components [48, 49]. For this reason, the AIC likely does not contribute to motor planning proper. Activity in the AIC can be more likely attributed to metacognition and a “feeling of knowing” (e.g. of a correct decision or of a memory item) [48, 49]. Accordingly, in the context of our study insular activity could refer to a “feeling of knowing” the relevant targets and/or movement plans.

Importantly, while we found broad activation across multiple brain areas during multi-target planning, we show that neural correlates of prospective motor planning in humans are concentrated in areas previously related to motor planning, but not in control areas V1l and M1l. We compare the activity modulation in all aforementioned areas based on a normative information theoretic bounded rationality model, where we find the highest explanatory power in terms of *R*^2^ for area SPL. Hence, our results support a leading role of human SPL in prospective parallel planning of target-directed movement sequences.

It would be clearly interesting to further investigate whether or not our network of planning-ROIs would differentially contribute to delay period activity during prospective parallel planning vs. serial processing (i.e. target memory). While our study was not designed in a way that would allow for such analyses, preliminary results of an earlier multivariate decoding analyses do not support such assumption [50].

### Limitations in memory and planning

In our study we have investigated limitations of concurrent planning in terms of the number and the sophistication of multiple plans. Limitations in parallel motor performance have been previously studied in dual-task designs. Considering the cognitive costs of dual task performance, serial processing frameworks [51, 52] were originally proposed that aim to explain delayed reaction times for secondary tasks [53]. Accordingly, the Response Selection Bottleneck (RSB) model [54] assumes that the crucial limitation (or bottleneck) in dual-task performance is located in the response selection stage. In contrast, capacity sharing models [55, 56] allow for sharing of cognitive resources in the central processing stages of two tasks. By manipulating task features like stimulus onset asynchrony and no-go trials, it has also been argued that subjects can somewhat switch between serial and parallel processing strategies [57, 58, 59, 60, 61]. As demonstrated in the current study, our information-theoretic model assumes constrained information processing for action planning and could be adapted to either delayed planning or concurrent prospective planning scenario, to allow for a quantitative analysis.

Another important limitation is imperfect memory of the stimulus during action planning. The concept of working memory as a short-term active storage of information [62] relates to the idea of a limited capacity control process involved in keeping or discarding information [63]. Quantifying the limited information capacity of working memory has been addressed in numerous studies, for example on visual working memory [19, 21, 22, 23, 64, 24] and spatial working memory [26, 27, 65, 66] with growing evidence that they share a common resource limitation [67, 68]. In terms of neurophysiology, it has been suggested early on, that memory limitations can be considered as the result of the limited computational capacity of single neurons [69]. In our model, we have represented limitations of working memory more abstractly, by an internal variable that can only assume a finite amount of values and whose precision is curtailed by a corresponding information constraint. This abstraction is based on the idea that any information processing limitation can ultimately be thought of as a limitation in the amount of uncertainty that can be reduced [70].

### Information theory, behavior and neural signals

Our study belongs to a large family of investigations that have used information theory to quantitatively model information processing constraints in behavior and neural signals. In behavioral experiments information capacity limits have been established for working memory, attention, perceptual identification, decision-making, motor planning and execution [19, 71, 72, 29]. Informational complexity in behavioral tasks is often related to reaction or movement times and task accuracy. The most well-known examples of this are Hick’s law that relates reaction time linearly to the entropy of a choice stimulus and Fitts’ law that relates movement time to an index of difficulty measured in bits [73, 74]. However, similar approaches have also been used to estimate the capacity for cognitive control depending on the entropy of the stimulus [75]. In our task, reaction times and error rates increased with higher task complexity, as expected from previous studies on motor sequence learning [3]. However, directly relating this change in reaction time to information complexity of our two hypotheses is confounded, because it is not clear whether the increase in reaction time during the response phase stems from a more complex decision between different ready-made motor plans (H_1_) or whether the increase results from finishing a single motor plan based on the incomplete uncertainty reduction during the delay phase (H_0_).

Importantly then, by going beyond the behavioral analysis, we could use the different information profiles for concurrent and serial planning to regress the fMRI signal. However, simply applying information-theoretic concepts to brain signals without taking into account the behavioral task, would not have been enough to distinguish between the two hypotheses. Previous neurophysiological studies have, for example, used mutual information between stimuli/behavior and neural signals to establish a neural code either for encoding or decoding [76], quantified the richness of conscious states within a theory of integrated information [77], or related activity in distinct regions of the prefrontal cortex to sensorimotor and cognitive control [78, 79]. A considerable number of imaging studies have also found neural correlates of informational surprise in sensory areas of the brain [80, 81, 82] related to prediction error in the context of predictive coding theory, with a wide range of applications ranging from action understanding in the mirror neuron system [83, 84, 85] and value-based decision-making [86] to hierarchical motor control [87]. However, in order to go beyond information-theoretic quantification of neural signals we need to consider both information constraints and task-related constraints [88].

### Bounded rationality

In contrast to many previous studies that have relied on information theory to estimate neural processing capacity, we have employed a class of bounded rationality models that trade off both utility and information processing costs [89, 90, 91, 92, 93], similar to the rate distortion trade-off in information theory [94] that defines relevant information when bandwidth is limited. Such a generalized trade-off makes this class of models applicable to tasks with arbitrary utility function. Moreover, using information constraints on multiple variables, we can design optimality models that simultaneously explain behavior and optimal information flow between internal variables that can be correlated with fMRI BOLD signals. In previous behavioral studies, the trade-off between utility and information has been used in the context of rational inattention and quantal response equilibria in economic decision-making [95, 96], perceptual representation in identification and memory tasks [72, 97], reaction time and endpoint variability for motor planning under time constraints [29], abstraction in sensorimotor processing [30], decision-making by sampling [98] and planning in a Markov environment [99, 100].

Similar to these previous studies where behavioral task performance and information processing capacities were measured, we here use a normative probabilistic optimality model for uncertainty reduction and concurrent processing. In this study, however, we go a step further and relate the predicted information flow with fMRI activity during motor planning. In particular, measuring the activity of premotor and parietal planning areas during the delayed response tasks allowed us to measure planning capacity for movement preparation. As the bounded rationality framework allows for multiple information constraints including internal variables for memory and action planning, we could regress fMRI activity with respect to these different information quantities and we could test the hypothesis of concurrent prospective planning (**H**_1_) against the null hypothesis of mere uncertainty reduction (**H**_0_). This approach rests on the assumption that brain signals reflect at least approximately an optimal information flow - an assumption that has been applied very successfully in the past, for example in the context of sparse coding [101, 102]. Ultimately, it is an empirical question for future studies, how far this class of models can be developed in their explanatory power, but they open a new exciting avenue that allows relating both behavior and neural representations to optimal information processing assumptions.

## 4 Acknowledgments

This study was funded by European Research Council (BRISC: Bounded Rationality in Sensorimotor Coordination).

## 5 Author contributions

S.S., A.L. and D.A.B. designed the experiment; S.S. and A.L. developed the experimental setup and acquired data; S.S., D.A.B. generated theoretical predictions from computer simulations; A.L., D.A.B supervised the project; S.S., A.L. and D.A.B. wrote, discussed and edited the manuscript.

## 7 Methods

### Experimental Methods

#### Subjects

Nineteen right-handed healthy subjects (11 females, 8 males) in the age range of 22 and 41 years (mean = 27.5, SD = 4.5) participated in the experiment and were included in further data analysis. Initially, we recruited 22 subjects, but data of three subjects was excluded because of strong movement artifacts. Participants provided written informed consent in accordance with the declaration of Helsinki and the study was approved by the ethics committee of the University Hospital and the Faculty of Medicine at the University of Töbingen.

#### Experimental Paradigm

We employed a delayed response task (DRT), in which subjects were instructed to generate sequences of key presses with four possible keys to control right, left, up, and down of a cursor movement. Each trial started with a random fixation phase of 13.5 - 16 sec (FIX), where a fixation cross was presented in the middle of the four fields and should be fixated with the eyes. This fixation period was followed by three principle task phases: cue presentation, delay/planning and response. In the cue presentation phase, subjects were shown one or multiple potential targets (four different target planning conditions *c_T_* ∈ {‘1’, ‘11’, ‘2’, ‘4’} as detailed below) with one of three possible sequence lengths (three different step conditions *c_S_* ∈ {2, 3, 4}). The timeline of a DRT is illustrated in Figure 1.

##### Cue presentation phase

The display area is divided into four panels each made up of 3×5 fields as shown in Figure 1 A). Each field represents a target position that can be reached with a cursor. Panels could be surrounded by a frame, which indicated that the highlighted target position within the frame should be taken into consideration as a potential target for motor planning. In all trials (including the training phase), only target positions with a sequence length of 2, 3 or 4 were considered due to the step conditions *c_S_* ∈ {2, 3, 4} of the experimental design. The potential target positions were always shown for 3 sec (CUE). Depending on the target planning condition, there were four different kinds of cues:

- ‘1’-Target Planning Condition: In this one-fold planning condition only one panel was framed containing a single highlighted potential target location that always corresponded to the actual target location in the response phase.
- ‘11’-Target Planning Condition: In this two-fold planning condition there was also a single framed panel that contained two potential targets highlighted, only one of which was selected as the actual target during response phase.
- ‘2’-Target Planning Condition: In this two-fold planning condition, two panels were framed, each containing a single highlighted potential target from which one was selected for execution.
- ‘4’-Target Planning Condition: In this four-fold planning condition all four panels were framed, each containing a single potential target highlighted for planning, from which one was selected in the response phase.

Cue presentation was followed by a mask of 1 sec (MASK), which was intended to “over-write” any after-images of the cue.

##### Planning phase

The planning phase was a delay phase of 14-16.5 sec (DEL) following the mask to allow subjects to plan one or multiple movement paths based on the information they obtained from the visual target cues. During this time, only a fixation cross was displayed in the center of the screen. From trial start to the completion of the planning phase, subjects had to keep their finger in rest position and where therefore asked to press the central button of the input device.

##### Response phase

In the response phase (RES), the actual target was determined from the set of potential targets and a go-signal. This go-signal comprised a half-size-frame encompassing six fields (including the actual target) was displayed around one half of one of the four field panels, however, the correct target itself was not highlighted in any way. Accordingly, the correct target field could only be uniquely identified through this go-signal if subjects remembered the appropriate information from the previously displayed target cues. Subjects were instructed to perform the movement sequence towards the inferred target field as fast and directly as possible within the allowed time window of 3 sec (otherwise the trial was terminated with an incomplete response). Depending on the distance of the target (sequence length), only the first 2, 3 or 4 key presses were considered before the trial was terminated. Subjects could observe the corresponding cursor movement and got immediate visual feedback of their performance at the end of the trial, by highlighting the cursor position in green (red) when the final reached position was correct (incorrect).

Besides DRT trials, we also included control trials (CT) in which no planning was possible during the delay phase. Like the other DRT trials, control trials were comprised an initial fixation phase and the three principal task phases – cue presentation, planning and response. Different to the DRT, however, in the cue presentation phase, no targets were highlighted. Hence, subjects were not informed about any potential target location and therefore could not plan specific movements during the delay planning phase. In the response phase, the actual target was indicated directly by highlighting the corresponding target field, so that the movement sequence to reach the target could only be planned once the actual target location was revealed.

Our experiment usually consisted of four blocks of 30 trials each (one subject had five blocks, two subjects only three blocks) typically resulting in a total of 120 trials. The overall experiment lasted ~ 80 min per subject. Each block consisted of 6 control trials and 6 trials for each target planning condition. Moreover, the 6 trials had an equal share of step conditions, i.e. 2 trials with sequence length of 2, 3, and 4, respectively. The order of the trials in any given block was permuted randomly.

In our experimental setup we used a Windows 7 based computer with R2007b (The MathWorks, Inc.) and the Cogent Graphics Toolbox (developed by John Romaya at the LON at the Wellcome Department of Imaging Neuroscience) to generate the visual stimuli for the experimental trial sequences and to record and store subjects’ behavioral responses and all relevant information associated with the visual stimuli sequence. The visual stimuli were back-projected by an LCD projector (1024×768 pixels; 60 Hz refresh rate) onto a semi-transparent screen at the end of the scanner bore behind the head coil. Subjects viewed the screen via a mirror that was mounted on top of the head coil. Screen dimensions amounted to 28 deg x 37 deg visual angle.

#### Behavioral performance monitoring

The instructed finger movements were monitored using the 5 Button Diamond Fibre Optic response device (Current Design, Philadelphia, US) operated by repetitive thumb flexions. From this we could extract reaction and movement times as well as error rates defined by the frequency of target misses.

For all behavioral analysis, we excluded trials, when movements were not initiated (1%) to compute reaction times and error rates and trials, when movements were not initiated or completed (2%), to compute movement times.

Additionally, eye movements were monitored at 50 Hz sampling rate throughout the scanning sessions using an MRI-compatible eye camara and infra-red illumination system (SMI SensoMotoric Instruments) and Viewpoint Eye Tracker software (Arrington Research, Scottsdale, US). Eye position recordings were filtered using a 10 Hz Chebyshev Type II low-pass filter. Saccades were detected using an absolute velocity threshold of 30 °/s (for a single recording of one subject 40 °/s). Eye data from 2 subjects were excluded from the analysis because of incompleteness.

We performed 2-way repeated measures ANOVAs (rmANOVA) with the factor target condition (‘1’, ‘11’, ‘2’, ‘4’ and CT) and factor sequence length (2, 3, or 4) for the behavioral analysis of reaction times, movement times, error rates and saccades. In addition we applied pairwise post-hoc comparisons of the respective marginal means. For this and all other rmANOVAs we tested for sphericity (Mauchly’s test) and applied the Greenhouse-Geisser correction whenever the assumption of sphericity was violated.

#### MRI image acquisition and analysis

MRI images were acquired on a 3 T Siemens PRISMA scanner (Siemens, Ellwangen, Germany). A T1-weighted magnetization prepared rapid-acquisition gradient echo (MP-RAGE) structural scan (176 slices, 256×256 voxel resolution, 1×1 mm inplane voxel size, slice thickness = 1 mm, gap = 0 mm, repetition time = 2300 ms, echo time = 2.96 ms, field of view = 256 mm) was obtained in parallel to the task training. Functional T2*-weighted gradient-echo planar imaging (EPI) volumes (48 sclices, 64×64 voxel resolution, 3×3 mm inplane voxel size, slice thickness =3 mm, gap = 0 mm, repetition time = 2000 ms, echo time = 35 ms, field of view = 1344 mm, flip angle=75 deg) completely covered the cerebellar cortex as well as subcortical structures. Per subject we obtained 600 EPIs during a single session of 19.75 min length, resulting overall in 2400 EPIs for 16 of the subjects (1800 and 2177 for prematurely terminated recordings and 3000 for a subject with an extra-session). Functional image processing was conducted with the SPM12 software package (Wellcome Centre for Human Neuroimaging, London, UK, Matlab 2016a Release) and included spatial realignment of all functional images to the first EPI image as a reference, co-registration of the anatomical T1 image and the mean functional EPI image, spatial normalization to the Montreal Neurological Institute space (MNI-template) with voxel size of 1×1×1 mm^3^ for T1 and 3×3×3 mm^3^ for EPI images and spatial smoothing of the normalized EPIs with a Gaussian kernel (7 mm full-width at half maximum (FWHM) Gaussian filter.

Functional fMRI analysis was first performed on an individual level (first-level analysis) and then on the group level (second-level analysis). The subject-specific analysis of fMRI BOLD activity is based on a general linear model (GLM). To this end for each session we specified 5 boxcar regressors for the cue phase of 3 sec (CUE) for each of the four target planning conditions (*c_T_*) and the control condition, as well as 5×3 boxcar regressors both for the delay phase of 14-16.5 sec (DEL) and for the response phase of 3 sec (RES), namely for each of the four target planning conditions (*c_T_*) and the control condition and separately for each of the three step conditions (*c_S_*). All aforementioned boxcar regressors were convolved with the canonical hemodynamic response function in SPM12. In addition, the 6 motion parameters from the realignment procedure were additionally included as regressors of no interest. The fixation phase was not explicitly modeled and served as an implicit baseline. Each experimental session therefore was modelled separately with 41 regressors for each subject (5 × CUE + (5*3)xDEL + (5*3) × RES + 6× motion regressors).

Regions of interest (ROIs) were selected as brain areas that exhibited significant planning-related activity in the DRT conditions compared to the control-task across subjects. Given the estimated regressor parameters of the GLM, in the 2nd-level analysis, we first determined significant voxels whose t-values for the contrast of interest (DRT_*Delay*_ > CT_*Delay*_) were below a threshold of *p* < 0.05 (t-test across subjects’ first level contrast images for DRT_*Delay*_ > CT_*Delay*_ family-wise error FWE-corrected for multiple comparisons). We found significant bilateral activities in brain areas typically involved in motor planning and visual memory [18]: posterior parietal cortex (superior parietal lobule (SPL) and anterior intraparietal sulcus (antIPS)), dorsal premotor cortex (PMd), dorsolateral prefrontal cortex (DLPFC), anterior insular cortex (AIC), cerebellum in lobulus VI and VIII (cer6 and cer8) and supplementary motor area (SMA). As a group coordinate of a ROI, we selected those voxel coordinates that were characterized by a maximum t value in the group analysis. Accordingly, we defined 15 group coordinates for the aforementioned areas (note that for the SMA, functional activity could not reliably be associated with either left or right hemisphere and therefore we only defined a single ROI). Also, we included left primary motor cortex (M1l) due to its contribution for our task related finger movements of pressing a button and therefore to control for activity related to movement execution directly. Similarly, to control for stimulus related activity we included left primary visual cortex (V1l). We identified the two anatomical regions of primary visual cortex (V1l) and primary motor cortex (M1l) with the corresponding group contrasts (CUE>0) and (RES–CUE; this was to account for the visual information present in the response phase).

For each individual subject, we mapped group ROI coordinates to individual ROI coordinates. This was done for each ROI, namely by identifying the coordinates of the local statistical maximum in the subject-specific first-level contrast (DRT_*Delay*_ > CT_*Delay*_) that was closest to the group-coordinates.

Group MNI coordinates of all 17 defined ROIs together with the average spatial dispersion of the ROI center on the individual subjects can be found in Supplementary Table 9.

For each subject and ROI, we extracted the GLM mean model parameters for the regressors of interest from a 3 mm radius sphere around each individual ROI coordinate. These beta parameters were (session-wise) normalized to the residual beta for any given session (i.e. baseline) to provide an estimate of the %-signal change of the fMRI signal. We computed rmANOVAs over fMRI beta estimates with factor target planning condition (‘1’, ‘11’, ‘2’, ‘4’) and with factor sequence length (2, 3, 4). In addition, event-related time courses of the fMRI-signal time courses (ERTs) of signal intensities were extracted using NERT4SPM [18].

### Theoretical Methods

#### Bounded Rationality Model

We applied the information-theoretic bounded rationality framework [28, 103] to our experimental task, where information about the hidden world state 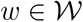 (the exact target location) is revealed in two steps, first by the initial cue stimulus 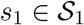 indicating all possible target locations, and after a delay by the go-signal 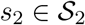 that resolves the remaining uncertainty about the actual hidden target *w*. After seeing the cue stimulus, the agent can form a memory 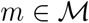 during the planning delay phase, before perceiving the cue stimulus *s*_2_ and subsequently selecting an action 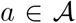 that corresponds to a movement sequence leading to a target location. The sets 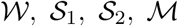 and 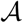 are finite and discrete, with 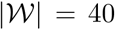 possible target locations (in distance of 2,3 or 4 steps), 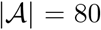 possible movement sequences, 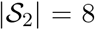 possible half-sized frames for each panel and 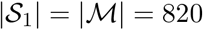 possible combinations of potential targets over all conditions. For any particular target condition *c_T_* and step condition *c_S_*, we then get a uniform distribution over stimuli

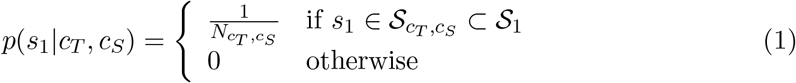

where 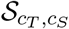 is the subset of stimuli that belong to the condition (*c_T_, s_T_*) and 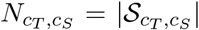 is the number of stimuli in that condition, where *N*_1,2_ = 8, *N*_1,3_ = 16, *N*_1,4_ = 16, *N*_11,2_ = 4, *N*_11,3_ = 16, *N*_11,4_ = 16, *N*_2,2_ = 24, *N*_2,3_ = 96, *N*_2,4_ = 96, *N*_4,3_ = 16, *N*_4_, 3 = 256, and *N*_4,4_ = 256. The marginal distribution over stimuli is given by *p*(*s*_1_) = ∑_*c_T_,c_S_*_*p*(*c_t_, c_S_*)*p*(*s*_1_|*c_T_, c_S_*) where all conditions are equally likely with 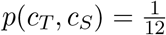. As the second stimulus *s*_2_ surrounds any potential target from *s*_1_ with a rectangular half-frame, the distribution *p*(*s*_2_|*s*_1_) is a uniform with non-zero probability over a single value of *s*_2_ when *c_T_* =‘1’, two possible values of *s*_2_ when *c_T_* =‘11’ or *c_T_* =‘2’, or four possible values of *s*_2_ when *c_T_* =‘4’.

In the model, the agent chooses the action a to maximize the task utility *U*(*w, a*) under the constraint that only a certain amount of information-processing can be achieved when forming the memory m and selecting the action *a*. For our task the utility function *U* is a simple 0/1-utility: it is 1 whenever the action *a* is compatible with the hidden target *w* and 0 otherwise.

Since information-processing can be unreliable, the agent’s memory state and action policy are formalized by probability distributions *p*(*m*|*s*_1_) and *p*(*a*|*m, s*_2_), such that the amount of information processing can be captured by the Kullback-Leibler divergence between the prior distributions *p*(*m*) and *p*(*a*) and the posterior distributions *p*(*m*|*s*_1_) and *p*(*a*|*m, s*_2_), respectively. The bounded rational decision-making problem can then be written as a constrained optimization problem

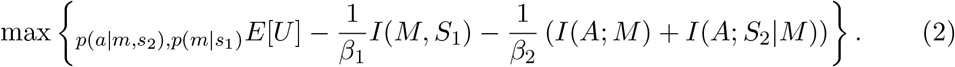

where the expectation is taken with respect to the distribution

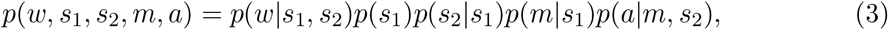

with *p*(*s*_1_), *p*(*s*_2_|*s*_1_) and *p*(*w*|*s*_1_, *s*_2_) defined by the task, and *p*(*m*|*s*_1_) and *p*(*a*|*m, s*_2_) left for optimization. Accordingly, the expected utility is given by

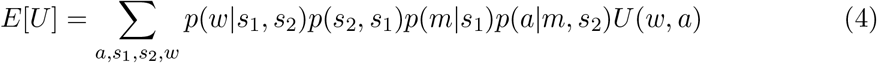

where *U*(*w,a*) = 1 if the action a leads to a target hit with target *w* and *U*(*w,a*) = 0 otherwise. Thus, we have *E*[*U*] = 1 – *ErrorRate*.

The information quantities *I*(*M, S*_1_) and *I*(*A*; *M*) + *I*(*A*; *S*_2_|*M*) respectively measure the average Kullback-Leibler divergence between the distributions *p*(*m*) and *p*(*m*|*s*1) for memory formation and the average Kullback-Leibler divergence between the distributions *p*(*a*) and *p*(*a*|*m, s*_2_) for generating a specific action given memory m and stimulus *s*_2_. The parameters *β*_1_ and *β*_2_ reflect the degree of boundedness, where *β*_1,2_ → ∞ reproduces a Bayes-optimal maximum expected utility decision-maker. The bounded optimal solution for equation 2 is given by

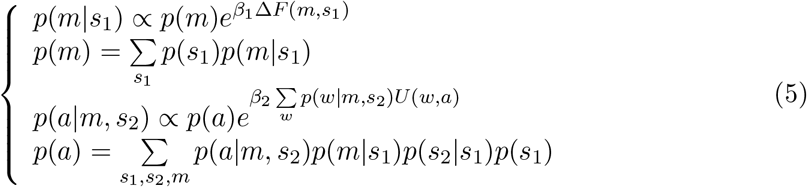

with

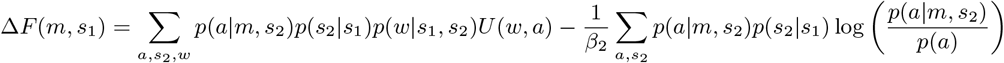

and with p(*w*|*m, s*_2_) determined from equation 3.

#### Modelling information-processing during Delay Phase

If we want to apply equation (5) to model planning during the delay phase, we have to be aware that the decision rule *p*(*a*|*m, s*_2_) requires the agent to know the go-stimulus *s*_2_ when deciding about the action a, an information that is not available during the delay period. We consider two hypotheses for information-processing during the delay phase.

- Hypothesis 0: Delayed Planning. Planning of the action *a* according to *p*(*a*|*m, s*_2_) is delayed until the response phase, once *s*_2_ is known. The delay phase is only used for uncertainty reduction, modeled by the mathematical transition from the prior *p*(*m*) to the posterior *p*(*m*|*s*_1_) and in the space of actions from *p*(*a*) to *p*(*a*|*m*). In any particular condition (*c_T_, c_S_*), we would then expect the information costs

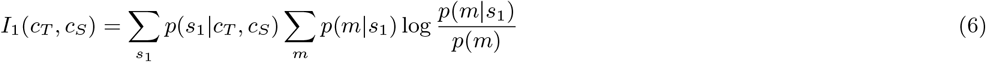

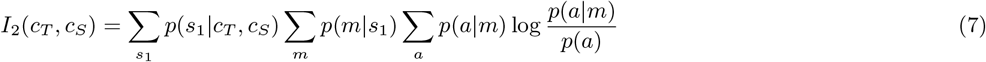

for memory formation and action processing respectively, and the average over all conditions given by

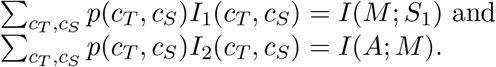
- Hypothesis 1: Prospective Planning. Once the uncertainty over actions is reduced to *p*(*a*|*m*) by observing *s*_1_, all possible *s*_2_ are anticipated in the delay phase, and for each *s*_2_ an action is planned according to *p*(*a*|*m, s*_2_). Depending on available information resources, the plans can be more or less precise. Once the actual go-stimulus *s*_2_ is revealed during the response phase, one of the planned actions can be immediately carried out. During the delay phase in any particular condition (*c_T_,c_S_*), we would then expect the information costs

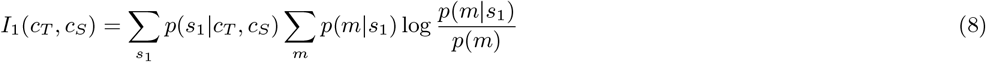

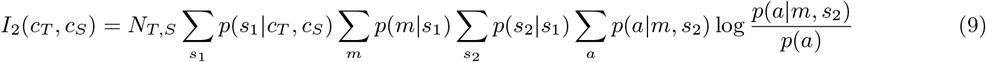

for memory formation and action processing respectively, with *N*_1,*S*_ = 1^2^, *N*_11,*S*_ = 2^2^, *N*_2,*s*_ = 2^2^ and *N*_4,*S*_ = 4^2^, and the average over all conditions given by

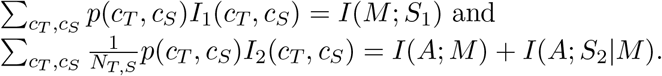

Assuming a linear relationship between informational surprise and brain signal fMRI(*c_T_, c_S_*) for each condition (*c_T_, c_S_*), we get a linear regression model

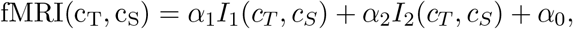

with model parameters *α_i_* = 0, 1, 2.

For each of the two hypothesis H_0_ and H_1_, we tested the multilinear regression between the fMRI activity modulation and the two information modulations predicted by the models, and with a nested model F-statistic (*α* = 5%), to find if *I*_2_ significantly improves the regression. Therefore, we applied the statistical test to the models with subject’s individual best fitting capacities and for individual fMRI activities in all relevant ROIs.

In a supplementary analysis, we repeated the regression for three different non-linearity assumptions regarding the relation between BOLD activity and information, namely quadratic

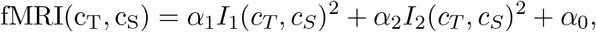

logarithmic

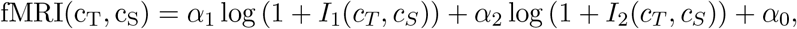

or sigmoidal

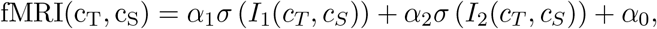

where 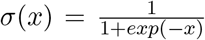. This allows to draw conclusions on the robustness of our hypothesis test results (see Supplementary Figure S4 and Supplementary Table 1).

