## Supplementary Material for "Bounded rational decision-making models suggest capacity-limited concurrent motor planning in human posterior parietal and frontal cortex"

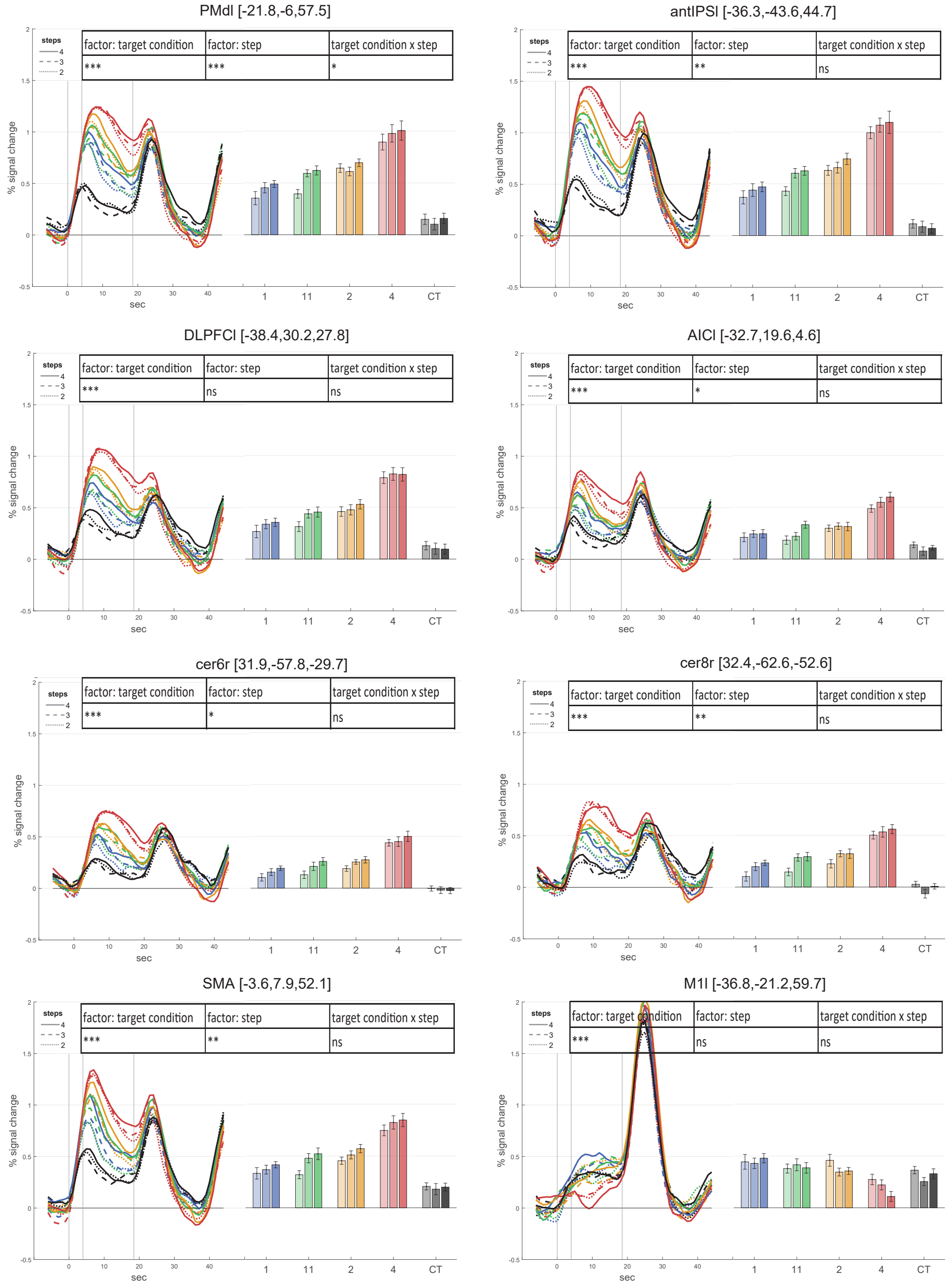


Supplementary Figure S 1: fMRI time course for different ROIs. Time course of raw activity as percentage of signal change normalized to pre-stimulus baseline and delay-activity estimates (beta parameter from GLM) in each planning condition in planning-related areas (PMdl, antIPSl, DLPFCl, AICl, cer6r, cer8r) show increasing BOLD amplitude with increasing task complexity. Primary left motor cortex (M1l) as a control area shows no significant difference between conditions. We report across-subjects averages and within-subjects variance as the normalized standard error (according to [31]) and average MNI-coordinates (x, y, z in mm) for each ROI. Statistical results are indicated with *** for $p <= 0.001$, ** for $0.001 < p <= 0.01$, * for $0.01 < p < 0.05$, and ns for non-significant results ($p > 0.05$).

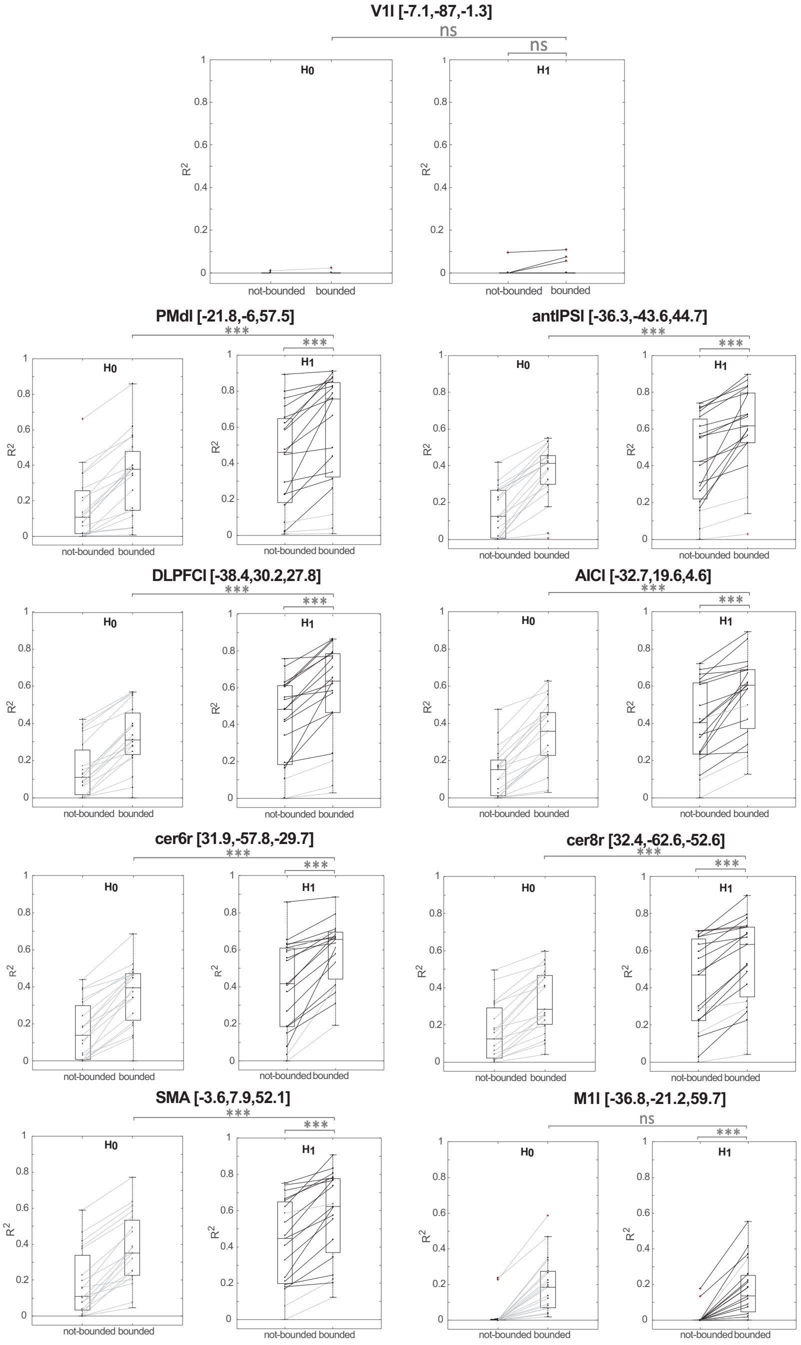


Supplementary Figure S 2: Model comparison for different ROIs. Similar to SPll, in all other planning related ROIs (PMdl, antIPSl, DLPFCl, AICl, cer6r, cer8r), a within-subject comparison of the $R^2$-values for the two hypotheses $H_0$ and $H_1$ showed that the information profiles under the prospective planning hypothesis $H_1$ provide a significantly better explanation of the fMRI modulation than the delayed planning hypothesis $H_0$ i ($p < 0.0001$ rmANOVA). The bounded rationality model predictions explain less of the fMRI activity modulation in control areas V1l and M1l with no significant difference between the model hypothesis ($p = 0.88$ and $p = 0.84$). We found that the correlation of measured brain activities in all other planning related ROIs (SPLl, PMdl, antIPSl, DLPFCl, AICl, cer6r, cer8r) and primary motor area (M1l) was significantly increased for model predictions with subject-individual bounds ($p < 0.0001$ rmANOVA) compared to an unbounded maximum capacity model. Only for control area V1l there was no significant difference between the bounded and unbounded model ($p = 0.131$). Statistical results of rmANOVAs are indicated with *** for $p <= 0.001$ and ns for non-significant results ($p > 0.05$).

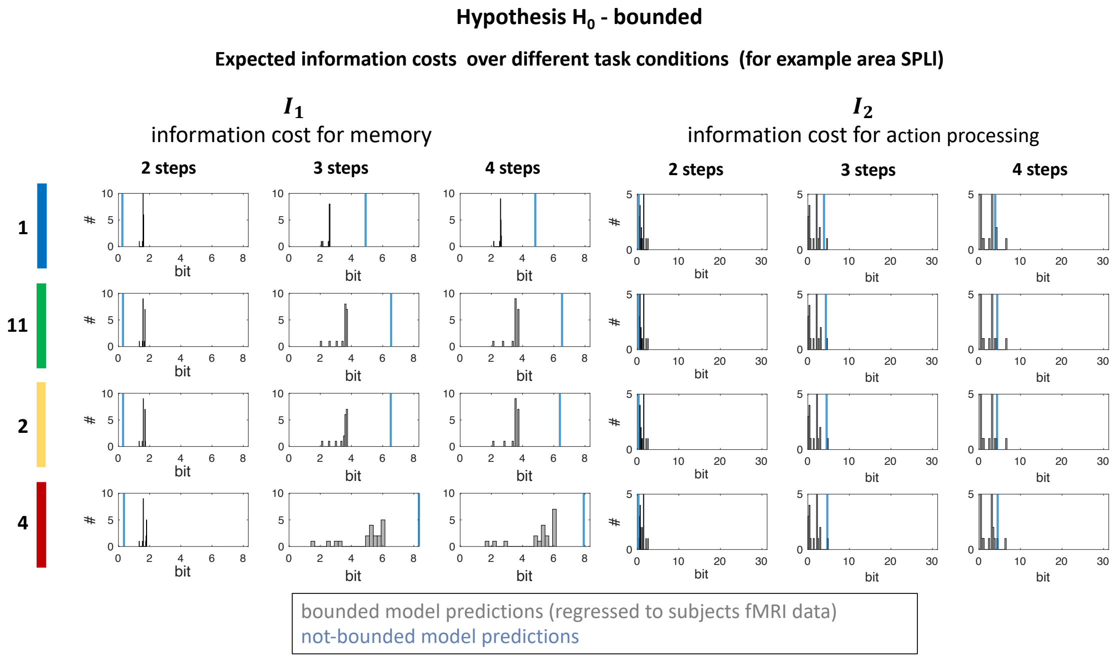


Supplementary Figure S 3: Expected information costs for hypothesis $H_0$ and $H_1$. Theoretical expected information costs $I_1$ (left) and $I_2$ (right) of the delayed planning hypothesis is based on model parameters determining memory and planning capacities fitted for all individual subjects (hypothesis $H_0$ "bounded"). Histograms represent the frequency distributions of the information costs, dependent on task conditions. Information values varied between bounded model predictions regressed to subjects fMRI data compared to the model predictions in the not-bounded case (in blue). Compared to the parallel planning hypothesis $H_1$, information $I_2$ for action processing is lower and planning postponed until the $s_2$-stimulus reveals the actual target location in the response phase of the experiment.

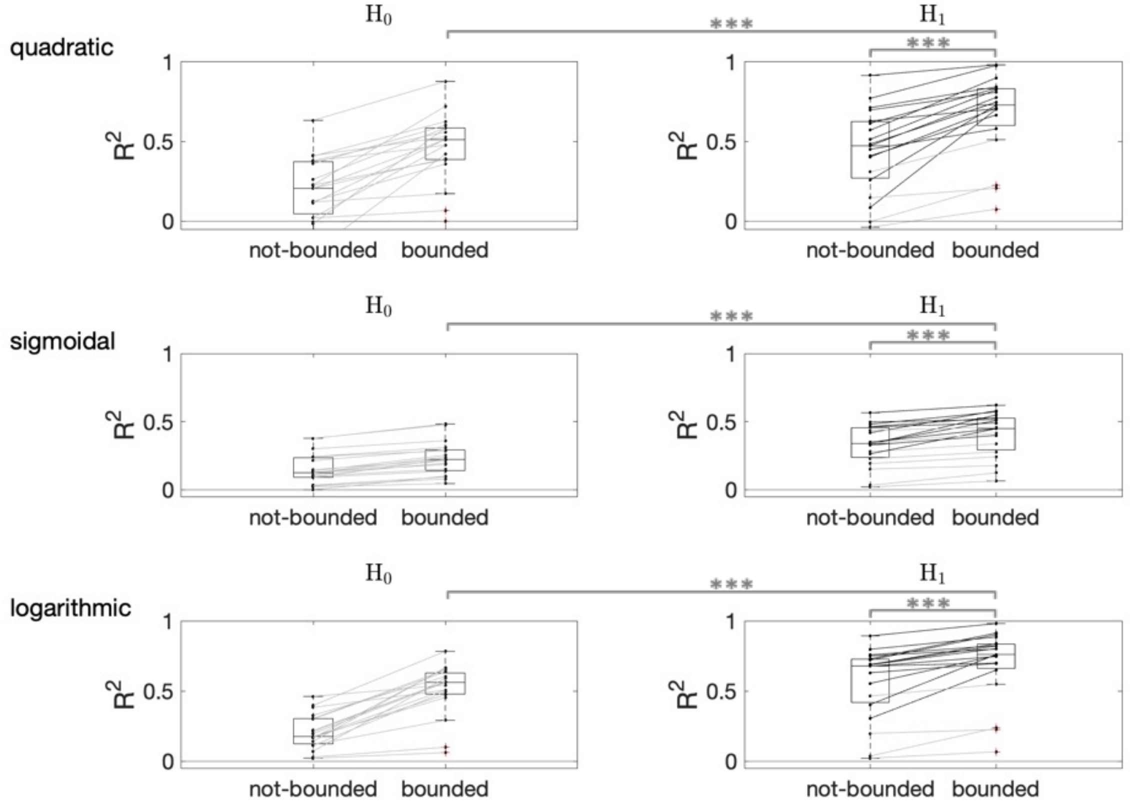


Supplementary Figure S 4: Model comparison for non-linear assumptions. Comparison of different non-linear assumptions (quadratic, sigmoidal, logarithmic) between fMRI correlates and predicted information values for model fits of example area SPLl. Similarly to the linear assumption (compare Figure 5), $R^2$-values of the regression analysis of the bounded rationality model under the prospective planning hypothesis $H_1$ are significantly higher than for the delayed planning hypothesis $H_0$ for the regressions of measured brain activities in SPLl as well as in all other planning related ROIs (also compare Supplementary Table 1). Likewise, compared to the not-bounded $H_1$ model hypothesis with maximum capacities ($H_1$ not-bounded), the bounded $H_1$ model hypothesis ($H_1$ bounded) with subject individual capacity fits could significantly better predict measured fMRI data in SPLl as well as in all other planning related ROIs (also compare Supplementary Table 1). Statistical results of rmANOVAs are indicated with *** for $p <= 0.001$.

Table 1: P-values for linear and non-linear assumptions. P-values of hypothesis tests $H_0$ vs $H_1$ and $H_1$ not-bounded (n-b) vs $H_1$ bounded (b) for linear and different non-linear relation assumptions (quadratic, sigmoidal, logarithmic) between fMRI correlates and predicted information values.

| | linear | | quadratic | | sigmoidal | | logarithmic | |
|---|---|---|---|---|---|---|---|---|
| | $H_0$ vs $H_1$ | $H_1 n-b$ vs $H_1 b$ | $H_0$ vs $H_1$ | $H_1 n-b$ vs $H_1 b$ | $H_0$ vs $H_1$ | $H_1 n-b$ vs $H_1 b$ | $H_0$ vs $H_1$ | $H_1 n-b$ vs $H_1 b$ |
| SPLl | 0.000 | 0.000 | 0.000 | 0.000 | 0.000 | 0.000 | 0.000 | 0.000 |
| PMdl | 0.000 | 0.000 | 0.000 | 0.000 | 0.000 | 0.000 | 0.001 | 0.000 |
| DLPFCl | 0.000 | 0.000 | 0.000 | 0.000 | 0.000 | 0.000 | 0.005 | 0.000 |
| antIPSl | 0.000 | 0.000 | 0.000 | 0.000 | 0.000 | 0.000 | 0.002 | 0.000 |
| AICl | 0.000 | 0.000 | 0.000 | 0.029 | 0.000 | 0.000 | 0.133 | 0.000 |
| cer6r | 0.000 | 0.000 | 0.000 | 0.001 | 0.000 | 0.000 | 0.000 | 0.000 |
| cer8r | 0.000 | 0.000 | 0.000 | 0.069 | 0.000 | 0.000 | 0.016 | 0.000 |
| SMA | 0.000 | 0.000 | 0.000 | 0.000 | 0.000 | 0.000 | 0.020 | 0.000 |
| V1 | 0.088 | 0.131 | 0.047 | 0.109 | 0.597 | 0.000 | 0.023 | 0.000 |
| M1 | 0.084 | 0.000 | 0.077 | 0.007 | 0.004 | 0.000 | 0.671 | 0.000 |

Table 2: $R^2$-values for hypothesis $H_0$ and $H_1$. $R^2$-values of the regression analysis of the bounded rationality model under the hypothesis $\mathbf{H}_0$ and $\mathbf{H}_1$. Group median with lower and upper quartiles of the bounded model assumptions (best fitting information capacities of individual subjects) are compared to the not-bounded case.

| | $H_0$ - not-bounded | | | $H_1$ - not-bounded | | |
|---|---|---|---|---|---|---|
| | median | 75% quantile | 25% quantile | median | 75% quantile | 25% quantile |
| SPLl | 0.12 | 0.28 | 0.04 | 0.51 | 0.64 | 0.34 |
| PMdl | 0.12 | 0.27 | 0.01 | 0.46 | 0.65 | 0.18 |
| DLPFCl | 0.10 | 0.25 | 0.01 | 0.48 | 0.61 | 0.18 |
| antIPSl | 0.11 | 0.26 | 0.01 | 0.42 | 0.65 | 0.22 |
| AICl | 0.15 | 0.21 | 0.01 | 0.40 | 0.62 | 0.23 |
| cer6r | 0.13 | 0.31 | 0.01 | 0.42 | 0.61 | 0.19 |
| cer8r | 0.12 | 0.28 | 0.02 | 0.47 | 0.66 | 0.22 |
| SMA | 0.09 | 0.32 | 0.04 | 0.45 | 0.65 | 0.20 |
| V1 | 0.00 | 0.00 | 0.00 | 0.00 | 0.00 | 0.00 |
| M1 | 0.00 | 0.00 | 0.00 | 0.00 | 0.00 | 0.00 |

| | $H_0$ - bounded | | | $H_1$ - bounded | | |
|---|---|---|---|---|---|---|
| | median | 75% quantile | 25% quantile | median | 75% quantile | 25% quantile |
| SPLl | 0.41 | 0.52 | 0.28 | 0.74 | 0.83 | 0.58 |
| PMdl | 0.38 | 0.48 | 0.15 | 0.76 | 0.85 | 0.32 |
| DLPFCl | 0.31 | 0.46 | 0.23 | 0.64 | 0.78 | 0.47 |
| antIPSl | 0.41 | 0.45 | 0.30 | 0.62 | 0.79 | 0.52 |
| AICl | 0.36 | 0.46 | 0.23 | 0.60 | 0.69 | 0.37 |
| cer6r | 0.39 | 0.47 | 0.22 | 0.66 | 0.70 | 0.44 |
| cer8r | 0.28 | 0.47 | 0.20 | 0.63 | 0.73 | 0.35 |
| SMA | 0.35 | 0.53 | 0.23 | 0.62 | 0.78 | 0.37 |
| V1 | 0.00 | 0.00 | 0.00 | 0.00 | 0.00 | 0.00 |
| M1 | 0.19 | 0.27 | 0.07 | 0.14 | 0.25 | 0.05 |

Table 3: $H_0$ bounded Hypothesis. Expected information $E[I_1]$ over all experimental conditions for all 19 subjects, measured in bits. For maximal capacity, information $E[I_1] = 4.428$ bits.

| subjects | $SPLl$ | $PMdl$ | $DLPFCl$ | $antIPS$ | $AICl$ | $cer6r$ | $cer8r$ | $SMA$ | $V1l$ | $M1l$ |
|---|---|---|---|---|---|---|---|---|---|---|
| 1 | 3.24 | 3.24 | 3.06 | 3.24 | 3.08 | 3.25 | 3.24 | 3.08 | 1.49 | 1.67 |
| 2 | 2.97 | 2.97 | 2.64 | 3 | 2.97 | 2.64 | 1.82 | 2.97 | 1.77 | 2.97 |
| 3 | 3.03 | 3.03 | 3.03 | 3.03 | 3.08 | 3.06 | 3.25 | 3.03 | 1.49 | 3.09 |
| 4 | 3.25 | 3.25 | 3.24 | 3.25 | 3.08 | 3.25 | 3.24 | 3.25 | 4.62 | 1.67 |
| 5 | 3.08 | 3.08 | 3.25 | 3.25 | 3.25 | 3.25 | 3.25 | 2.64 | 1.77 | 2.97 |
| 6 | 3.03 | 3.03 | 1.38 | 1.38 | 1.38 | 1.38 | 2.97 | 1.38 | 1.43 | 1.49 |
| 7 | 3.24 | 3.09 | 3.24 | 3.11 | 3.09 | 3.08 | 3.23 | 3.03 | 1.53 | 1.51 |
| 8 | 3.03 | 3.06 | 3.06 | 3.08 | 3.08 | 3.08 | 3.24 | 3.09 | 1.77 | 3.09 |
| 9 | 1.77 | 2.78 | 1.77 | 1.72 | 3.09 | 1.73 | 1.72 | 3.09 | 1.77 | 1.74 |
| 10 | 2.65 | 3.25 | 3.16 | 3.16 | 3.17 | 2.65 | 3.22 | 3.15 | 1.96 | 2.97 |
| 11 | 3.25 | 3.03 | 3.18 | 3.25 | 3.24 | 3.21 | 3.25 | 3.18 | 1.77 | 1.96 |
| 12 | 3.09 | 3.06 | 3.06 | 3.1 | 3.25 | 2.43 | 3.08 | 3.08 | 1.77 | 1.82 |
| 13 | 3.03 | 3.09 | 3.03 | 3.03 | 3.06 | 3.03 | 3.25 | 3.03 | 1.77 | 1.8 |
| 14 | 2.3 | 2.78 | 3.19 | 3.12 | 3.09 | 1.54 | 3.12 | 3.09 | 1.54 | 3.09 |
| 15 | 3.24 | 3.19 | 3.19 | 3.24 | 1.67 | 1.8 | 3.17 | 1.67 | 1.77 | 1.67 |
| 16 | 3.24 | 3.24 | 3.24 | 3.24 | 3.24 | 3.25 | 3.24 | 3.18 | 1.54 | 1.72 |
| 17 | 3.24 | 3.24 | 3.24 | 3.24 | 3.24 | 3.24 | 3.24 | 3.24 | 1.77 | 1.7 |
| 18 | 2.97 | 3.09 | 2.97 | 2.97 | 3.07 | 3.24 | 3.08 | 2.97 | 1.77 | 1.72 |
| 19 | 2.62 | 2.82 | 2.62 | 2.62 | 2.88 | 2.5 | 2.24 | 2.82 | 1.77 | 2.97 |
| mean | 2.96 | 3.07 | 2.92 | 2.95 | 2.95 | 2.72 | 2.99 | 2.89 | 1.85 | 2.19 |

Table 4: $H_0$ bounded Hypothesis. Expected information $E[I_2]$ over all experimental conditions for all 19 subjects, measured in bits. For maximal capacity, information $E[I_2] = 2.994$ bits.

| subjects | $SPLl$ | $PMdl$ | $DLPFCl$ | $antIPS$ | $AICl$ | $cer6r$ | $cer8r$ | $SMA$ | $V1l$ | $M1l$ |
|---|---|---|---|---|---|---|---|---|---|---|
| 1 | 2.32 | 2.32 | 0.7 | 2.32 | 1.01 | 2.42 | 2.32 | 0.93 | 2.65 | 2.97 |
| 2 | 0.4 | 0.4 | 0.36 | 0.53 | 0.4 | 0.36 | 0.61 | 0.4 | 2.57 | 0.4 |
| 3 | 0.53 | 0.53 | 0.53 | 0.53 | 0.93 | 0.64 | 2.42 | 0.53 | 2.65 | 2.39 |
| 4 | 2.42 | 2.42 | 2.32 | 2.42 | 1.01 | 2.42 | 2.32 | 2.42 | 2.88 | 2.97 |
| 5 | 0.93 | 0.93 | 2.42 | 2.42 | 2.42 | 2.42 | 2.42 | 0.36 | 2.57 | 0.4 |
| 6 | 0.53 | 0.53 | 0.42 | 0.42 | 0.42 | 0.42 | 0.4 | 0.42 | 2.52 | 3.09 |
| 7 | 2.32 | 2.39 | 2.32 | 2.61 | 2.39 | 0.93 | 2.3 | 0.53 | 2.62 | 3.03 |
| 8 | 0.53 | 0.7 | 0.7 | 0.93 | 0.93 | 0.93 | 2.32 | 2.39 | 2.57 | 2.39 |
| 9 | 2.57 | 2.5 | 2.57 | 3.16 | 2.39 | 3.04 | 3.16 | 2.39 | 2.57 | 2.98 |
| 10 | 0.46 | 2.4 | 2.38 | 2.38 | 2.4 | 0.46 | 2.37 | 2.39 | 2.76 | 0.4 |
| 11 | 2.42 | 0.53 | 2.37 | 2.42 | 2.32 | 2.54 | 2.42 | 2.37 | 2.57 | 2.85 |
| 12 | 2.39 | 0.7 | 0.7 | 2.42 | 2.42 | 0.22 | 0.93 | 0.93 | 2.57 | 0.61 |
| 13 | 0.53 | 2.39 | 0.53 | 0.53 | 0.64 | 0.53 | 2.42 | 0.53 | 2.57 | 3.1 |
| 14 | 3.38 | 2.5 | 2.36 | 2.64 | 2.39 | 2.85 | 2.64 | 2.39 | 2.85 | 2.39 |
| 15 | 2.32 | 2.42 | 2.36 | 2.32 | 2.97 | 3.1 | 2.44 | 2.97 | 2.57 | 2.97 |
| 16 | 2.32 | 2.32 | 2.32 | 2.32 | 2.32 | 2.42 | 2.42 | 2.37 | 2.85 | 3.16 |
| 17 | 2.32 | 2.32 | 2.32 | 2.32 | 2.32 | 2.42 | 2.42 | 2.42 | 2.57 | 2.59 |
| 18 | 0.4 | 2.39 | 0.4 | 0.4 | 1.37 | 2.42 | 2.63 | 0.4 | 2.57 | 3.16 |
| 19 | 3.19 | 3.04 | 3.19 | 3.19 | 3.1 | 1.41 | 3.42 | 3.04 | 2.57 | 0.4 |
| mean | 1.7 | 1.77 | 1.65 | 1.91 | 1.8 | 1.68 | 2.23 | 1.59 | 2.63 | 2.22 |

Table 5: $H_1$ bounded Hypothesis. Expected information $E[I_1]$ over all experimental conditions for all 19 subjects, measured in bits. For maximal capacity, information $E[I_1] = 4.428$ bits.

| subjects | $SPLl$ | $PMdl$ | $DLPFCl$ | $antIPS$ | $AICl$ | $cer6r$ | $cer8r$ | $SMA$ | $V1l$ | $M1l$ |
|---|---|---|---|---|---|---|---|---|---|---|
| 1 | 2.86 | 1.74 | 2.6 | 1.69 | 1.67 | 3.14 | 2.44 | 2.36 | 1.49 | 1.8 |
| 2 | 2.78 | 2.78 | 2.73 | 2.97 | 2.97 | 1.82 | 1.82 | 2.78 | 1.77 | 1.82 |
| 3 | 3.08 | 3.07 | 3.08 | 3.09 | 3.1 | 3.06 | 3.09 | 2.82 | 2.12 | 1.49 |
| 4 | 3.2 | 3.14 | 3.1 | 3.17 | 3.18 | 3.12 | 3.1 | 3.17 | 4.94 | 1.67 |
| 5 | 3.17 | 3.16 | 2.57 | 2.27 | 2.12 | 2.25 | 2.41 | 2.32 | 1.77 | 1.82 |
| 6 | 3.05 | 3.07 | 3.07 | 3.06 | 3.08 | 3.09 | 2.77 | 3.08 | 1.43 | 1.49 |
| 7 | 3.09 | 2.97 | 3.08 | 1.69 | 3.18 | 1.68 | 1.69 | 2.65 | 1.53 | 1.51 |
| 8 | 3.1 | 3.09 | 3.1 | 3.12 | 3.1 | 3.16 | 3.1 | 3.08 | 1.77 | 1.82 |
| 9 | 2.97 | 1.72 | 2.97 | 2.97 | 2.12 | 1.73 | 1.72 | 3.01 | 1.77 | 1.74 |
| 10 | 1.8 | 1.8 | 1.8 | 1.97 | 3.19 | 2.24 | 1.82 | 3.15 | 1.96 | 2.12 |
| 11 | 3.11 | 2.83 | 3.04 | 1.89 | 4.8 | 2.42 | 3.16 | 3.11 | 1.77 | 1.96 |
| 12 | 2.7 | 2.53 | 3.09 | 1.67 | 2.8 | 2.83 | 3.03 | 3.09 | 1.77 | 1.82 |
| 13 | 3.01 | 3.03 | 3.11 | 3.05 | 3.09 | 3.1 | 3.17 | 3.1 | 1.77 | 1.8 |
| 14 | 2.3 | 1.7 | 3.09 | 1.7 | 3.1 | 2.97 | 1.7 | 2.97 | 1.54 | 1.67 |
| 15 | 1.69 | 1.69 | 1.7 | 1.69 | 1.8 | 1.8 | 1.67 | 1.8 | 1.77 | 1.67 |
| 16 | 3.08 | 3.08 | 3.08 | 3.01 | 3.1 | 2.43 | 1.77 | 1.7 | 2.25 | 1.72 |
| 17 | 1.68 | 1.68 | 1.74 | 2.44 | 1.7 | 1.77 | 4.8 | 1.7 | 1.77 | 1.72 |
| 18 | 2.73 | 2.65 | 2.12 | 2.7 | 1.8 | 1.7 | 1.69 | 2.78 | 1.77 | 1.72 |
| 19 | 1.91 | 2.62 | 2.62 | 2.47 | 2.73 | 2.62 | 2.08 | 2.62 | 1.77 | 2.05 |
| mean | 2.7 | 2.54 | 2.72 | 2.45 | 2.77 | 2.47 | 2.48 | 2.7 | 1.93 | 1.76 |

Table 6: $H_1$ bounded Hypothesis. Expected information $E[I_2]$ over all experimental conditions for all 19 subjects, measured in bits. For maximal capacity, information $E[I_2] = 14.109$ bits.

| subjects | $SPLl$ | $PMdl$ | $DLPFCl$ | $antIPS$ | $AICl$ | $cer6r$ | $cer8r$ | $SMA$ | $V1l$ | $M1l$ |
|---|---|---|---|---|---|---|---|---|---|---|
| 1 | 12 | 12 | 11.92 | 12.16 | 12.71 | 12.66 | 12.17 | 12.01 | 12.75 | 13.65 |
| 2 | 11.36 | 11.36 | 12.19 | 10.02 | 3.01 | 4.77 | 4.77 | 11.36 | 12.46 | 4.77 |
| 3 | 9.96 | 7.65 | 6.59 | 6.93 | 12.19 | 8.02 | 12.12 | 11.82 | 1.85 | 12.75 |
| 4 | 12.25 | 11.9 | 11.9 | 11.99 | 12.17 | 11.84 | 11.9 | 11.93 | 14.95 | 12.71 |
| 5 | 12.27 | 12.25 | 12.22 | 12.01 | 12.62 | 14.43 | 12.12 | 12.42 | 12.46 | 4.77 |
| 6 | 6.03 | 6.35 | 4.96 | 6.43 | 7.27 | 12.12 | 8.87 | 6.59 | 12.45 | 13.62 |
| 7 | 11.76 | 3.01 | 11.86 | 12.3 | 11.86 | 12.09 | 12.16 | 11.79 | 12.64 | 13.47 |
| 8 | 12.11 | 11.76 | 11.96 | 11.83 | 11.9 | 12.1 | 11.85 | 11.86 | 12.46 | 4.77 |
| 9 | 3.01 | 13.79 | 3.01 | 3.01 | 1.85 | 13.49 | 13.79 | 7.46 | 12.46 | 13.38 |
| 10 | 13.4 | 13.4 | 13.4 | 14.34 | 12.26 | 14.24 | 12.31 | 12.15 | 12.89 | 1.85 |
| 11 | 12.68 | 12.05 | 13.66 | 12.9 | 14.64 | 12.07 | 12.28 | 13.8 | 12.46 | 13.1 |
| 12 | 11.81 | 11.79 | 9.87 | 12.71 | 11.83 | 11.95 | 10.92 | 9.25 | 12.46 | 4.77 |
| 13 | 5.04 | 3.65 | 12 | 9.58 | 8.69 | 12.11 | 12.04 | 12.11 | 12.46 | 13.65 |
| 14 | 14.12 | 12.57 | 11.76 | 12.57 | 11.96 | 3.01 | 12.57 | 3.01 | 13.11 | 12.71 |
| 15 | 12.16 | 12.16 | 12.57 | 12.13 | 13.65 | 13.65 | 12.71 | 13.65 | 12.46 | 12.71 |
| 16 | 11.86 | 11.86 | 11.86 | 11.97 | 11.85 | 12.08 | 12.9 | 12.57 | 5.04 | 13.79 |
| 17 | 12.17 | 12.17 | 13.38 | 12.17 | 12.57 | 12.9 | 14.64 | 12.57 | 12.46 | 13.79 |
| 18 | 12.19 | 3.48 | 1.85 | 12.11 | 13.65 | 12.57 | 12.3 | 11.36 | 12.46 | 13.79 |
| 19 | 11.91 | 12.98 | 13.9 | 13.9 | 12.19 | 12.98 | 5.11 | 12.98 | 12.46 | 2.31 |
| mean | 10.95 | 10.32 | 10.57 | 11.11 | 10.99 | 11.53 | 11.45 | 11.09 | 11.73 | 10.33 |

Table 7: $H_0$ bounded Hypothesis. Expected utility $E[U]$ over all experimental conditions for all 19 subjects.

| subjects | $SPLl$ | $PMdl$ | $DLPFCl$ | $antIPS$ | $AICl$ | $cer6r$ | $cer8r$ | $SMA$ | $V1l$ | $M1l$ |
|---|---|---|---|---|---|---|---|---|---|---|
| 1 | 1 | 1 | 0.56 | 1 | 0.65 | 1 | 1 | 0.63 | 0.81 | 0.75 |
| 2 | 0.43 | 0.43 | 0.43 | 0.48 | 0.43 | 0.49 | 0.37 | 0.43 | 0.86 | 0.43 |
| 3 | 0.48 | 0.48 | 0.48 | 0.48 | 0.63 | 1 | 0.48 | 0.48 | 0.81 | 1 |
| 4 | 1 | 1 | 1 | 1 | 0.65 | 1 | 1 | 1 | 1 | 0.75 |
| 5 | 0.63 | 0.63 | 1 | 1 | 1 | 1 | 1 | 0.43 | 0.86 | 0.43 |
| 6 | 0.48 | 0.48 | 0.48 | 0.48 | 0.48 | 0.48 | 0.48 | 0.48 | 0.79 | 0.8 |
| 7 | 1 | 1 | 1 | 0.99 | 1 | 0.67 | 0.43 | 0.48 | 0.82 | 0.8 |
| 8 | 0.48 | 0.56 | 0.56 | 0.63 | 0.63 | 1 | 1 | 1 | 0.86 | 1 |
| 9 | 0.86 | 0.87 | 0.86 | 0.83 | 1 | 0.85 | 0.48 | 1 | 0.86 | 0.84 |
| 10 | 0.47 | 1 | 1 | 1 | 1 | 0.37 | 0.56 | 1 | 0.89 | 0.43 |
| 11 | 1 | 0.48 | 1 | 1 | 1 | 1 | 1 | 1 | 0.86 | 0.88 |
| 12 | 1 | 0.56 | 0.56 | 1 | 1 | 0.56 | 0.48 | 0.63 | 0.86 | 0.54 |
| 13 | 0.48 | 1 | 0.48 | 0.48 | 0.53 | 1 | 0.53 | 0.48 | 0.86 | 0.86 |
| 14 | 0.91 | 0.87 | 1 | 1 | 1 | 1 | 1 | 1 | 0.82 | 1 |
| 15 | 1 | 1 | 1 | 1 | 0.75 | 0.85 | 0.75 | 0.75 | 0.86 | 0.75 |
| 16 | 1 | 1 | 1 | 1 | 1 | 1 | 1 | 1 | 0.82 | 0.83 |
| 17 | 1 | 1 | 1 | 1 | 1 | 1 | 1 | 1 | 0.86 | 0.85 |
| 18 | 0.43 | 1 | 0.43 | 0.43 | 0.76 | 1 | 0.43 | 0.43 | 0.86 | 0.83 |
| 19 | 0.95 | 0.97 | 0.95 | 0.95 | 0.98 | 0.99 | 0.59 | 0.97 | 0.86 | 0.43 |
| mean | 0.77 | 0.81 | 0.78 | 0.83 | 0.81 | 0.86 | 0.71 | 0.75 | 0.86 | 0.75 |

Table 8: $H_1$ bounded Hypothesis. Expected utility $E[U]$ over all experimental conditions for all 19 subjects.

| subjects | $SPLl$ | $PMdl$ | $DLPFCl$ | $antIPS$ | $AICl$ | $cer6r$ | $cer8r$ | $SMA$ | $V1l$ | $M1l$ |
|---|---|---|---|---|---|---|---|---|---|---|
| 1 | 0.98 | 0.86 | 0.96 | 0.85 | 0.75 | 1 | 1 | 0.95 | 0.81 | 0.86 |
| 2 | 0.87 | 0.87 | 0.87 | 0.85 | 0.43 | 0.49 | 0.79 | 0.87 | 0.86 | 0.54 |
| 3 | 0.89 | 0.76 | 0.7 | 0.72 | 1 | 0.87 | 0.65 | 0.96 | 0.33 | 0.81 |
| 4 | 1 | 1 | 1 | 1 | 1 | 1 | 1 | 1 | 0.99 | 0.75 |
| 5 | 1 | 1 | 0.96 | 0.94 | 0.91 | 0.94 | 0.96 | 0.94 | 0.86 | 0.54 |
| 6 | 0.65 | 0.68 | 0.58 | 0.67 | 0.74 | 0.56 | 0.48 | 0.7 | 0.79 | 0.8 |
| 7 | 1 | 0.43 | 1 | 0.85 | 1 | 0.86 | 0.75 | 0.97 | 0.82 | 0.8 |
| 8 | 1 | 1 | 1 | 1 | 1 | 1 | 1 | 1 | 0.86 | 0.54 |
| 9 | 0.43 | 0.83 | 0.43 | 0.43 | 0.33 | 0.84 | 0.81 | 0.73 | 0.86 | 0.84 |
| 10 | 0.86 | 0.86 | 0.86 | 0.88 | 1 | 0.9 | 0.9 | 1 | 0.89 | 0.33 |
| 11 | 0.99 | 0.99 | 0.99 | 0.88 | 0.99 | 1 | 0.97 | 0.99 | 0.86 | 0.88 |
| 12 | 0.97 | 0.96 | 0.88 | 0.75 | 0.97 | 0.53 | 0.53 | 0.85 | 0.86 | 0.54 |
| 13 | 0.58 | 0.48 | 1 | 0.86 | 0.82 | 0.96 | 0.84 | 1 | 0.86 | 0.86 |
| 14 | 0.91 | 0.85 | 1 | 0.85 | 1 | 0.85 | 1 | 0.43 | 0.82 | 0.75 |
| 15 | 0.85 | 0.85 | 0.85 | 0.85 | 0.86 | 0.86 | 0.54 | 0.86 | 0.86 | 0.75 |
| 16 | 1 | 1 | 1 | 0.99 | 1 | 0.95 | 1 | 0.85 | 0.58 | 0.83 |
| 17 | 0.85 | 0.85 | 0.84 | 0.95 | 0.85 | 0.92 | 0.92 | 0.85 | 0.86 | 0.83 |
| 18 | 0.87 | 0.47 | 0.33 | 0.91 | 0.86 | 0.99 | 0.97 | 0.87 | 0.86 | 0.83 |
| 19 | 0.89 | 0.92 | 0.95 | 0.93 | 0.87 | 0.99 | 0.59 | 0.92 | 0.86 | 0.37 |
| mean | 0.87 | 0.82 | 0.85 | 0.85 | 0.86 | 0.87 | 0.83 | 0.88 | 0.82 | 0.71 |

Table 9: ROI coordinates. MNI coordinates of the group coordinates and average spatial dispersion of the ROI centers on individual subjects around the group ROI coordinates.

| | group coordinates | | | ± difference to individual coordinates | | |
|---|---|---|---|---|---|---|
| | $x$ | $y$ | $z$ | $x$ | $y$ | $z$ |
| SPLl | −9 | −69 | 57 | 6.63 | −2.68 | −0.63 |
| SPLr | 9 | −63 | 54 | −9 | 3.47 | −3.63 |
| PMdl | −18 | −3 | 63 | 3.79 | 3 | 5.53 |
| PMdr | 27 | −6 | 60 | 1.58 | −2.68 | 4.58 |
| DLPFCl | −36 | 30 | 24 | 2.37 | −0.16 | −3.79 |
| DLPFCr | 36 | 45 | 27 | −1.26 | 9.79 | −2.84 |
| antIPSl | −39 | −45 | 45 | −2.68 | −1.42 | 0.32 |
| antIPSr | 36 | −42 | 42 | −0.16 | −0.79 | −2.84 |
| AICl | −33 | 15 | 6 | −0.32 | −4.58 | 1.42 |
| AICr | 36 | 18 | 9 | 0.95 | −2.37 | 5.21 |
| SMA | 9 | 6 | 54 | 12.63 | −1.89 | 1.89 |
| V1l | 0 | −78 | 3 | 7.11 | 9 | 4.26 |
| M1l | −42 | −15 | 60 | −5.21 | 6.16 | 0.32 |
| cer6l | −27 | −57 | −33 | 1.89 | 1.26 | −2.68 |
| cer6r | 33 | −57 | −30 | 1.11 | 0.79 | −0.32 |
| cer8l | −33 | −60 | −48 | −0.63 | 1.58 | 4.89 |
| cer8r | 33 | −60 | −48 | 0.63 | 2.58 | 4.58 |

Table 10: Leave-one-out-cross-validation. Predictive ability of the suggested models was evaluated using leave-one-out cross-validation. Multi-linear regression of the measured fMRI modulation and the predicted information values $I_1$ and $I_2$ was performed on data with 11 out of 12 conditions in a repeated manner and regression parameters used to predict the fMRI activity in the left-out condition. In all brain areas except the control areas M1l and V1l, the prediction error under the concurrent prospective planning hypothesis $H_1$ is significantly lower than the prediction error under the delayed planning hypothesis H0. Similarly, under the prospective planning hypothesis $H_1$ the "bounded" models with subject-individual best-fitting information capacities ($H_1$ b) have a significantly lower prediction error in the cross-validation than the "not-bounded" models with maximum capacity($H_1$ n-b). Non-parametric pairwise comparison was performed using Wilcoxon signrank test with an Bonferroni corrected significance level ($p = 0.00294$), when testing for all the 17 ROIs (left and right hemispheres, see Methods).

| | $H_0$ vs $H_1$ | $H_1 n - b$ vs $H_1 b$ |
|---|---|---|
| SPLl | 0.00290 | 0.00290 |
| PMdl | 0.00097 | 0.00097 |
| DLPFCl | 0.00223 | 0.00223 |
| antIPSl | 0.00062 | 0.00062 |
| AICl | 0.00084 | 0.00084 |
| cer6r | 0.00016 | 0.00016 |
| cer8r | 0.00062 | 0.00062 |
| SMA | 0.00054 | 0.00054 |
| V1 | 0.25000 | 0.25000 |
| M1 | 0.04862 | 0.04862 |
